# Mycoelectronics: Bioprinted Living Fungal Bioelectronics for Artificial Sensation

**DOI:** 10.1101/2025.10.06.680560

**Authors:** Yulu Cai, Caleb Ronders, Vittorio Mottini, Hang Yuan, Kalpana Singh, Liuxi Xing, Iha Singh, Denghao Fu, Kyla Zhao, Linux Heller, Khoi Nguyen, Bryce Waller, Tuo Wang, Gregory Bonito, Jinxing Li

## Abstract

The intelligence of the human biological system is enabled by the highly distributed sensing receptors on soft skin that can distinguish various stimulations or environmental cues, thus establishing the fundamental logic of sensing and physiological regulation or response. To replicate biological perception, two approaches have emerged: artificial nervous systems that utilize soft electronics as biomimetic receptors to convert external stimuli into frequency-encoded signals, and biohybrid solutions that integrate living cells, plants, or even live animals with electronic components to decode environmental cues for life-like sensations. However, most current biohybrid approaches for artificial sensation are based on eukaryotic cells, which suffer from slow growth, stringent culture conditions, environmental susceptibility, and short lifespans, thus limiting their integration into practical wearables or robotic sensory skins. Here, we introduce fungi-based printable “*Mycoelectronics*”, which are created by additive bioprinting of living fungal mycelium networks onto stretchable electronics, as a practical living thermo-responsive sensory platform. This Mycoelectronics approach leverages fungi’s capacity for rapid biological responsiveness, cultivability with exponential growth, stability and self-healing in ambient conditions, bioprintability for scalable manufacturing, and mechanical flexibility for seamless integration with soft electronics. Critically, we discovered that the thermal responsiveness of the fungal network arises from intrinsic cellular processes—specifically, heat-induced vacuole remodeling and fusion, which modulate ionic transport and thus the electrical conductivity of the mycelial cells and networks, enabling a rapid temperature response. By bridging the gap between cell biology and soft electronics, the Mycoelectronics device with a living mycelium network functions as a thermal sensation system with rapid response and intrinsic self-healing properties, autonomously restoring sensing capabilities after damage or autonomously establishing sensor pathways in hard-to-reach locations. Furthermore, by integrating fungal thermal sensing with electronic circuits, we established a hybrid bioelectronic reflex arc that can actuate muscles and initiate diverse actions, suggesting promising applications in future neurorobotics and neuroprosthetics.

## Introduction

From swimming microbes to migrating cells, and ultimately to mammals and humans, the ability to sense and adapt to a changing environment is a fundamental aspect of life. Through evolution, living organisms have developed highly integrated architectures in which sensing, processing, and actuation are intertwined at both structural and functional levels(*1*). In humans, intelligence begins at the skin: a soft, compliant surface embedded with thousands of distributed sensory receptors capable of distinguishing subtle variations in pressure, vibration, and temperature(*2*). For example, the thermoreceptor is one of the most fundamental and ancient sensory mechanisms in biological systems(*3*, *4*). The adequate stimulus for a thermoreceptor is warming, which increases its action potential discharge rate(*5*). The temperature receptors innervate various organs, including the skin (as cutaneous receptors), viscera, and hypothalamus. This seamless coupling of structure and function, achieved through bottom-up morphogenesis and biological growth, enables organisms to maintain robust, adaptable, self-healing, and scalable sensory interfaces—capabilities that synthetic systems still struggle to replicate.

The biological sensation inspired the recent development of soft electronic skin and artificial sensation systems that can replicate the complex sensing capabilities of human skin and nerves for restoring sensation or enhancing robotic intelligence(*6–11*). In the past decade, a set of state-of-the-art artificial receptors(*12–14*), synapses(*15–18*), neurons(*19–21*), and nerves(*22–25*), has been developed. Enabled by the advanced materials engineering and design architecture, these soft artificial nervous systems have significantly advanced in signal processing and computing(*23*, *26–29*), self-powering(*30–32*), and incorporating multimodal sensation and responses(*13*, *33*, *34*). These artificial nervous systems have also been successfully applied to actuate biological muscles and control robotic systems(*24*, *25*, *35*). Despite notable success, their reliance on complex materials and fabrication processes often results in high costs and limited scalability(*8*, *36*). More critically, their efficiency, resilience, and versatility still fall short of natural biological systems, which are inherently regenerative, self-healing, and capable of seamlessly integrating multiple sensing modalities. This performance gap motivates the search for alternative, sustainable platforms that directly integrate living, adaptive biological networks to achieve robust, scalable, and multifunctional sensory capabilities.

Biohybrid devices offer a compelling strategy to address emerging challenges by integrating living biological components—such as living cells(*37–39*), microbes(*40–42*), muscle tissues(*43–45*), or even animals(*46*, *47*), plants(*48–50*)—with man-made electronic systems. For instance, cultured and genetically engineered rat cardiomyocytes grown on optoelectronic substrates have enabled the development of phototactically guided soft robotic rays that can sense and respond to light(*51*). Insects such as locusts and honeybees have been merged with electronics to create hybrid sensors capable of detecting volatile chemicals(*46*, *47*). More recently, microbial systems have been explored for environmental sensing(*40*, *52*), detection of biomarkers(*53*), and managing of inflammation(*42*). However, biohybrid systems based on eukaryotic cells still face significant limitations. Animal cell-based systems, such as those using skeletal muscles or cardiac muscle tissues(*44*), require tightly controlled sterile environments, daily medium exchange, and supplementation with nutrients and antibiotics to maintain viability. Moreover, plant-based systems often suffer from slow growth and the difficulty of directing morphogenesis, making them less practical for seamless integration with modern electronics(*54*). These challenges highlight the need for alternative biological platforms that are more scalable, resilient, and compatible with engineered systems.

Fungi are among the oldest eukaryotic life forms on earth, with mycelial networks that play critical roles in nutrient cycling and symbiotic support of plants—highlighting that environmental responsiveness and distributed intelligence can emerge from non-neural biological systems(*55–58*). They inhabit not only natural environments but also the human body(*59*, *60*). Research has shown that fungi can respond to environmental stimuli through various modes of complex communication(*56*, *61–64*). Environmental perturbations, such as heat stress(*65*), elicit rapid (second-level) and pronounced cellular responses that extend beyond gene expression. Compared to mammalian cells, plants, or bacteria, fungi form complex multicellular mycelial structures that offer mechanical integrity, electrical conductivity and biopotentials(*63*, *66*, *67*), and a hierarchical spatial organization that extends from individual hyphae to interconnected, large-scale subterranean webs(*68*), ideal for biohybrid interfaces. Moreover, owing to their intrinsic robustness, rapid proliferation, and environmental adaptability, fungi can be easily manipulated and processed in a wide range of conditions(*69–71*), significantly improving manufacturability, especially printability, for integration with synthetic devices.

By leveraging Mycological concepts with recent engineering advances in soft electronics and biomanufacturing, we present a new biohybrid electronics platform, bioprinted living fungal “Mycoelectronics” (Mycology + electronics), that uniquely integrates the resilience, printability, and sensing capabilities of living fungi with the precision of modern biofabrication. The Mycoelectronics system is assembled through a two-step printing process: first, laser-induced graphene (LIG) is patterned into soft electrode arrays; second, a bioink, made by living fungal mycelium and nutritive hydrogel, is directly and precisely printed onto these electrodes, forming a mechanically compliant, self-sustaining interface. This approach creates a structurally and functionally integrated thermo-responsive interface—biologically inspired, skin-mimicking, and scalable for biomanufacturing. By harnessing the mycelium’s natural growth-driven patterning, we enable tunable sensory networks capable of autonomous repair, navigating complex topographies, and large-area thermal sensing. The living Mycoelectronics self-assemble functional connections within hours and maintain stable operation for over 50 hours, remaining robust through daily tests for multiple days and retaining sensing capability after months of maintenance-free storage, with no need to replace the growth medium and only minimal recalibration before use. Integrating with closed-loop electronic circuits, this living network supports a fungal-based reflex arc for both sensing and actuation, laying the foundation for a new generation of self-healing, environmentally adaptive, and closed-loop devices for bioinspired robotics and neuroprosthetics.

## Results

Biological temperature perception is mediated by thermoreceptors located in free nerve endings, which transduce thermal stimuli into electrical signals that are transmitted through afferent nerves to the central nervous system for interpretation (Fig. 1A). Similarly, our fungal bioelectronic thermoreceptor operates at both cellular and network levels to transduce heat into measurable electrical changes. We discovered that at the cellular level, increased temperature induces vacuolar remodeling and fusion in fungal hyphae, characterized by an increase in vacuolar size. Due to the insulating nature of the vacuolar membranes, these changes physically obstruct ionic transport pathways, thereby modulating local conductivity (Fig. 1B). At the network level, such thermal stimulation collectively induces ionic redistribution across the interconnected hyphal network, leading to changes in the global conductivity of the mycelium. Together, these hierarchical responses form the basis of thermal sensing in living fungal electronics. The aerial mycelium network, therefore, as shown in Fig. 1C, functions as a thermal sensor by converting thermal stimuli into measurable changes in electrical conductivity through intimate connections to bottom graphene electrodes.

**Fig. 1.**
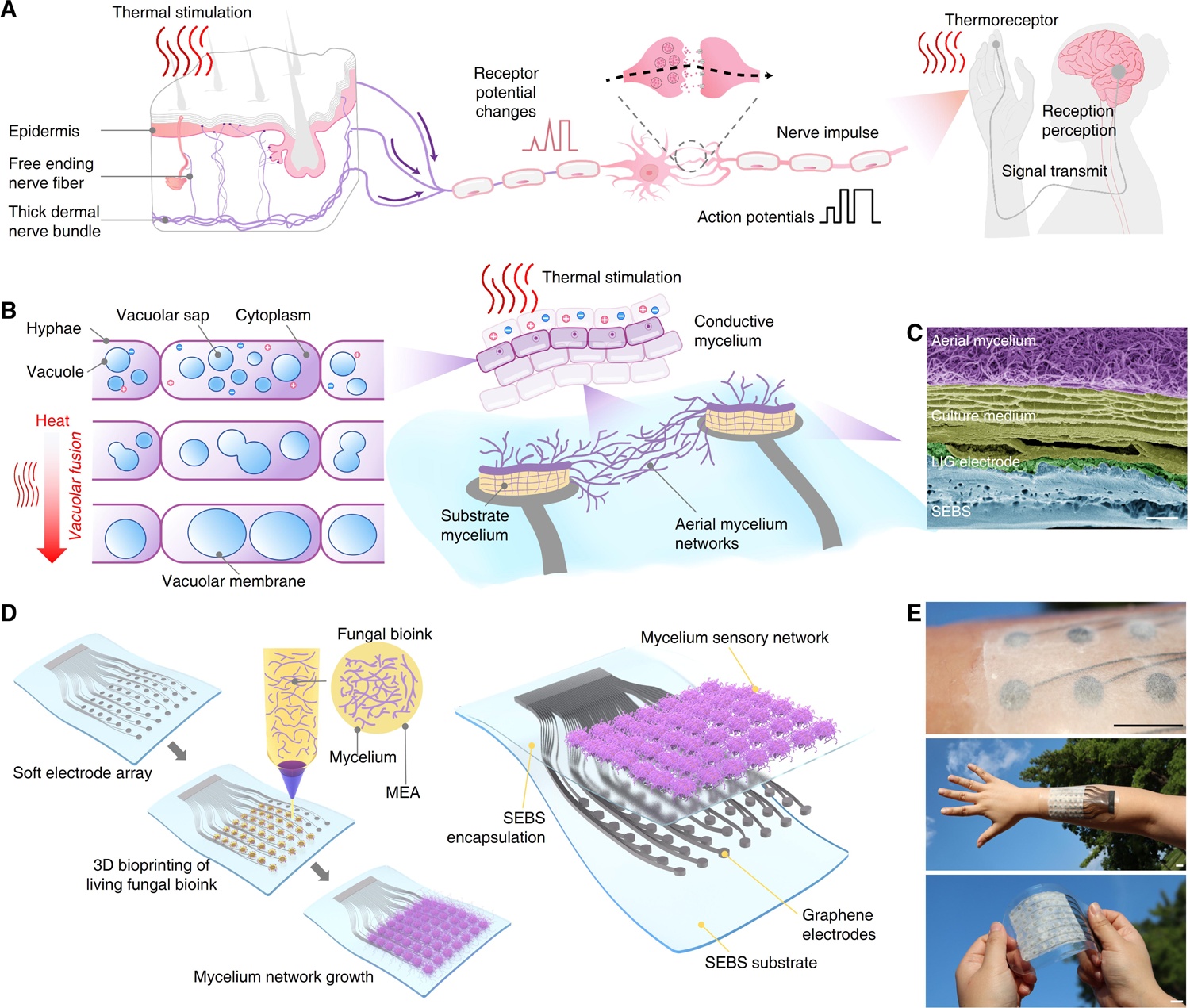
Living fungal Mycoelectronics thermoreceptor compared to a human thermoreceptor. **A**, Human thermoreceptor and sensory pathway from skin to brain. **B,** Mechanism of the Mycoelectronics thermoreceptor from the microscopic to the macroscopic scale: Thermal stimulation rapidly induces vacuolar fusion in fungal hyphae, transforming dispersed vacuoles into large compartments, the insulating vacuolar membranes occupy most of the hyphal volume and restrict ionic transport, thereby increasing electrical resistance. **C,** Scanning electron microscopy (SEM) image showing the microscale architecture of the electrode-mycelium interface of the 3D-bioprinted living Mycoelectronics (scale bar, 100 μm). **D,** Schematics showing the biological fabrication and growth of living fungal mycelium networks on a soft graphene-based electrode array to form the multiple-channel Mycoelectronics sensing array. **E,** Photographs show the mechanical flexibility and wearability of the 3D-bioprinted living fungal Mycoelectronics sensor array (scale bar, 1 cm).

The fungi and LIG planar electrodes are interfaced with a nutrient medium layer (using agarose or gelatin), cultivated with substratal mycelium in between, while aerial mycelium acts as a sensing bridge to connect the planar electrodes. Fig. 1D illustrates the biological printing and growth process for creating a multiple-channel Mycoelectronics sensing array. This system consists of a 3D bioprinted inoculated nutrient hydrogel on a stretchable electrode array, which is made by laser-patterned graphene electrodes embedded in a styrene-ethylene-butylene-styrene (SEBS) elastomer, a process developed in our previous work(*72*). After bioprinting, the hyphae are then grown in an ambient environment, forming a complex aerial mycelial network that bridges the electrodes and creates conductive pathways with thermo-responsive capabilities. This scalable and rapid biomanufacturing approach, along with self-sustaining growth in an ambient environment, represents a significant step toward the practical implementation of soft living sensing technologies (Fig. 1E). We observe that the graphene electrodes can form intimate contact with the growing mycelium, allowing a stable connection (Supplementary fig. S1-7 and Movie 1). For the fabrication of stretchable Mycoelectronics thermoreceptors, a gelatin-based culture medium was used. Gelatin’s elasticity and strong adhesion to the LIG electrode significantly stabilized the mycelial network structure (Supplementary fig. S8,9 and Movie 2). Our electrical stretchability test indicates a restoration of the initial impedance levels after 25% stretching, affirming the potential of Mycoelectronics as a soft and conformable bioelectronics platform (fig. S10 and Movie 3).

To create the biohybrid system, we first studied cultivation nutrients and media for controlled mycelial growth and bioprinting, which are necessary for developing a printable gel-based “bioink” with inoculated live mycelium for biomanufacturing. Unlike other biohybrid systems that utilize mammalian cells or bacteria, fungi exhibit robust and rapid growth across a wide range of nutrient levels. Here, *Benniella eriona* (a member of the *Mortierellaceae*, a commonly encountered non-pathogenic hyphae) is screened as the model fungus(*73*). We first evaluated the growth kinetics of *Benniella erionia* at varying concentrations of malt extract (ME) agar media (0%-10%) (Fig. 2A). *Benniella erionia* exhibits relatively rapid growth, achieving the highest radial expansion rate, 6 hours for 1 cm, at 2% ME concentration (Fig. 2B, C). However, higher ME concentrations suppressed colony expansion, likely due to osmotic stress or metabolic feedback regulation(*74*). To evaluate the mycelium’s bridging capability, we prepared cylindrical fungal samples (1 cm^2^) together with their underlying agar medium (3 mm thick) after one week of growth at different ME concentrations. These were transferred onto empty Petri dishes with predefined gap distances and monitored over seven days (Fig. 2D). Remarkably, the fungal network retained its bridging capability across gaps as large as five times its original size (5 cm), even under low-nutrient conditions. Higher ME concentrations promoted long-distance bridging; under 8% ME, a 10 cm gap was completely bridged within four days, the fastest among all tested conditions. To translate this biological growth behavior into practical fabrication, we developed a printable fungal ink by blending substratal mycelium with crushed culture medium (ME=2%) into a gelatinous matrix. Rheological characterization showed that the mycelium-gel ink as bioink maintained shear-thinning behavior comparable to a printable hydrogel (2% ME agar medium), enabling smooth extrusion during printing (Fig. 2E). Further three-interval thixotropy tests (3ITT) and step–strain sweep measurements revealed that the viscoelastic properties of the bioink were maintained under oscillatory shear (Fig. 2F and G). Strain amplitude sweeps revealed similar storage modulus (*G′*) and loss modulus (*G″*) moduli between the bioink and hydrogel inks, indicating that incorporating substratal mycelium did not compromise mechanical integrity (Fig. 2H). Based on these favorable material properties, we implemented a direct bioprinting strategy to pattern the mycelium-gel bioink onto stretchable graphene-SEBS electrode arrays with precise spatial control. As shown in Fig. 2I, we demonstrated multiple electrode layouts (black) paired with corresponding fungal printing pattern designs (purple) and photographs showing their integrations. This versatile design-to-device printing workflow enables precise spatial alignment of the bioink with the underlying electrode arrays. Subsequent growth and proliferation of aerial mycelium bridged the electrode gaps, forming biological networks that established electrical pathways and enabled sensing functionality (Fig. 2I right column). A schematic illustration (Fig. 2J top) depicts the 3D bioprinting process, where aerial mycelium grows within the bioink-defined boundaries, bridging electrode gaps to form sensing nodes. The growth patterns can be visualized by UV light due to the autofluorescence of the mycelium networks, as shown in the printed Spartan logo (Fig. 2J). Time-lapse images and Movie captured the fast and controllable extrusion of ink into the 49-channel electrode arrays, forming an “MSU” (Fig. 2K and Movie 4). After bioprinting, mycelial proliferation occurs within the defined pattern constraints over subsequent days, with growth patterns visualized through fluorescent light (Fig. 2L). Moreover, the flexibility of the overall material system allows the Mycoelectronics sensing array to conform to various curved surfaces, such as a human hand as a living wearable device, as shown in Fig. 2L right. This growth-driven biological patterning and fabrication demonstrates the potential of Mycoelectronics as a dynamic system capable of adapting to complex device geometries.

**Fig. 2.**
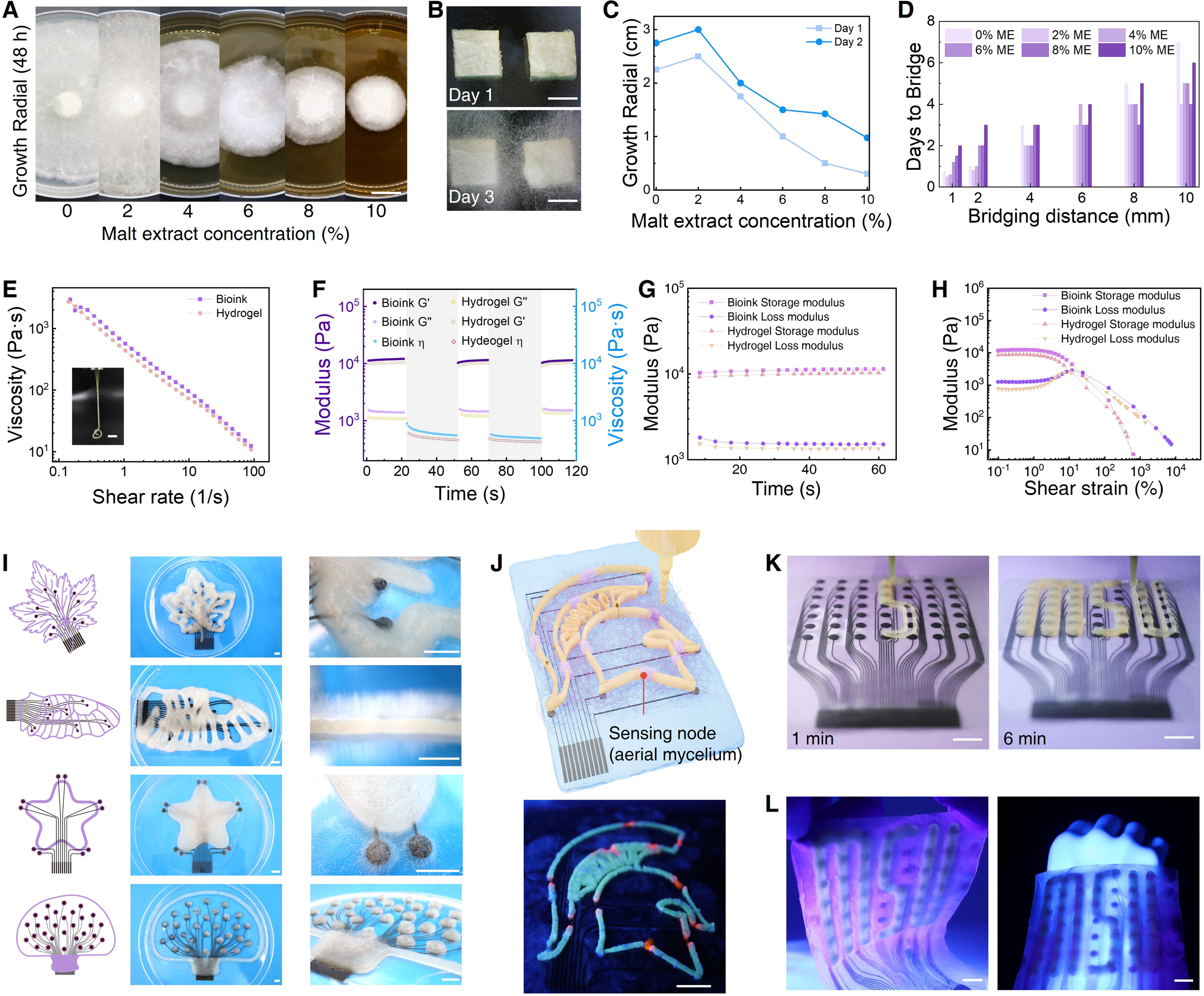
Growth behavior and rheological properties of Mycoelectronics inks, and bioprinting of fungal Mycoelectronics sensing devices. **A**, Photographs of fungal colony morphology after 48 h of growth on ME agar media at varying concentrations (0–10% w/v) (scale bar, 1 cm). **B,** Optical images showing aerial mycelial bridge between two electrodes from Day 1 (top) to Day 3 (bottom) (scale bar, 1 cm). **C,** Quantification of radial colony growth on Day 1 and Day 2 at different ME concentrations. **D,** Number of days required for mycelial bridging over gaps of 1–10 mm under different ME concentrations. **E,** Shear-thinning behavior of the bioink and hydrogel (scale bar, 1 cm). **F,** 3ITT profiles of the bioink and hydrogel after inducing structural disruption. **G,** Step–strain sweep confirming structural recovery of the bioink after large strain deformation. **H,** Nonlinear viscoelastic behavior and yield points through strain amplitude sweeps for the bioink and hydrogel. **I,** Schematic of the 3D bioprinting layout, where black lines represent LIG electrode arrays and purple paths indicate the printed mycelium–gel bioink. Representative printed patterns and corresponding optical close-ups are shown on the right (scale bars, 1cm). **J,** Schematic and photography of 3D bioprinting process in which an array with the living bioink printed as a “Spartan logo”. The lower image shows the fungal autofluorescence under UV light, where blue autofluorescence marks the printed bioink and red fluorescence (stained with Rhodamine B after growth) highlights mycelial growth bridging the gaps forming sensing nodes (scale bars, 1 cm). **K,** Time-lapse images of the fungal bioprinting process on a 49-electrode array at 1 min and 6 min (scale bars, 1 cm). **L,** Photographs of printed Mycoelectronics array after 5 days of growth (left) and demonstration of Mycoelectronics thermoreceptor’s flexibility by conformal attachment to a human hand (right) (scale bars, 1 cm).

The comprehensive characterization of the thermal response of the living sensors is first studied through planar electrodes (Fig. 3A and Movie 5). Sensory performance of the living Mycoelectronics was characterized by using a single-channel setup where thermal stimuli were added through a programmable voltage-controlled heating pad with a thermometer for calibration (Fig. 3B). The biological response was recorded through a designed single-channel setup that integrates graphene electrodes (black), culture medium (yellow) and the mycelium network (purple), as illustrated in Fig. 3C. Chronopotentiometry was employed to apply 1 nA current to the circuit, of which the mycelium penetrates the intermediate culture medium layer and directly contacts the LIG electrode, resulting in two double-layer capacitors (C_1_ and C_2_) between the LIG and both the mycelium and the medium. The ionic-conducting mycelium and medium comprise numerous parallel capacitors (CPE) and internal diffusion resistance (R_ct_). Since the current we applied will traverse the entire Mycoelectronics, the determining factor for the overall resistance changes is the aerial mycelium bridge. Impedance spectroscopy test (Fig. 3D) revealed the distinct impedance change at lower frequency range (<1 k Hz) under heating (red line) and subsequent return to room temperature (blue line). This region is related to *α* frequency range (corresponds to the ionic diffusion in cell membrane in the frequency range of 10 Hz to 10 kHz), as ions move across cellular barriers(*75*). Systematic thermal tolerance studies revealed that *Benniella erionia* maintains viable growth between 0-45 °C under both brief (1 min) and extended (10 min) heat exposures, while temperatures exceeding 55 °C slow the mycelial development (fig. S11), establishing a practical operating range for environmental sensing applications.

**Fig. 3.**
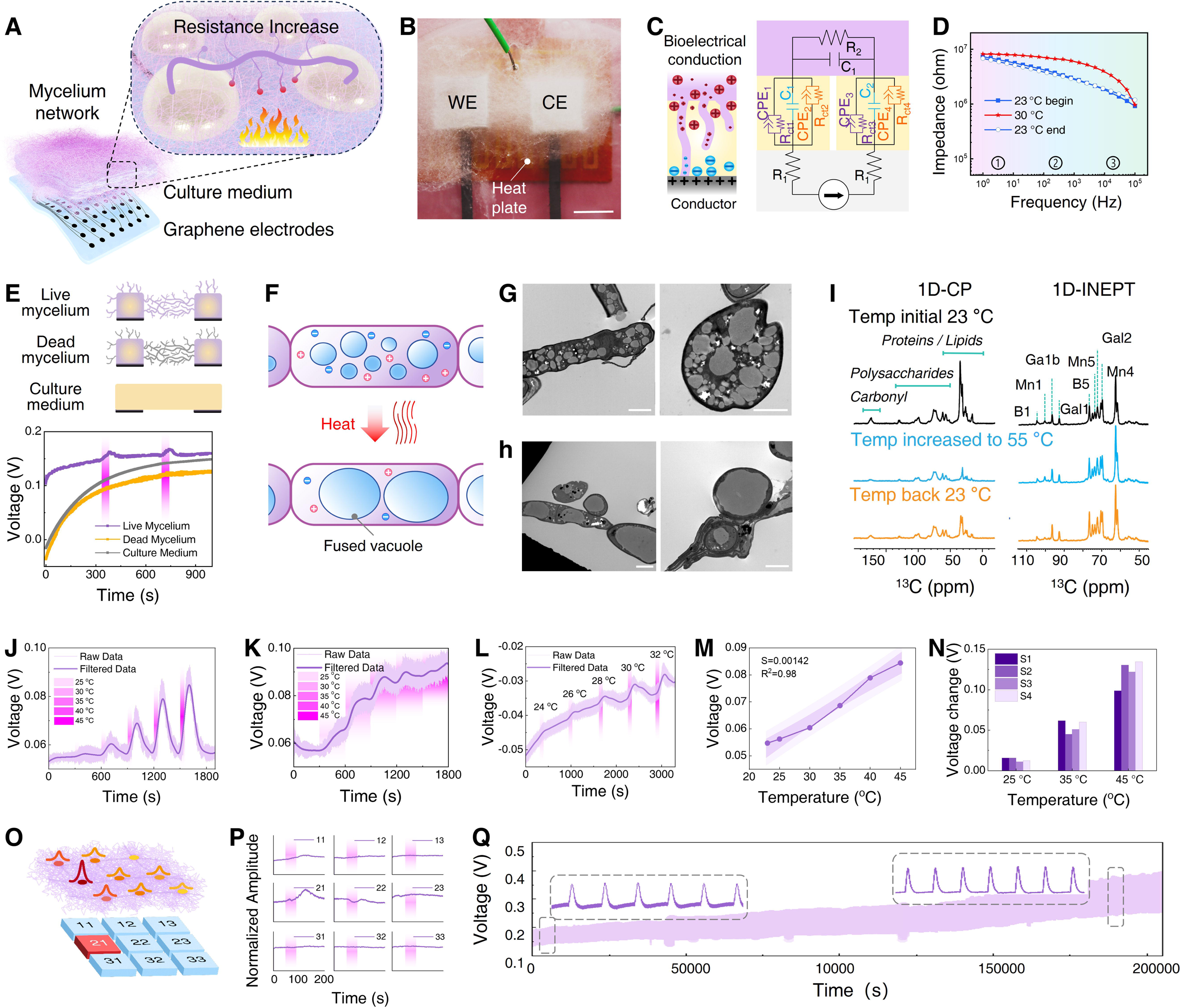
Characterization and mechanistic understanding of the Mycoelectronics thermoreceptor. **A**, Schematic showing increased mycelial resistance under heat exposure. **B,** Experimental setup utilizing a precision-controlled heating pad as the heat source, with thermometers for accurate temperature calibration (scale bar: 1 cm). **C,** Electrical circuit diagram depicting the interactions between graphene electrodes, culture medium, and mycelium network. **D,** Electrochemical impedance spectroscopy (EIS) test of single channel results showing a significant increase in low-frequency resistance with temperature circle (23 °C - 35 °C - 23 °C). **E,** Comparison of voltage responses between live mycelium, dead mycelium, and culture medium under a 65 °C, 10-min thermal stimulation. **F,** Schematic illustration of heat-induced vacuolar fusion within hypha. Upon heat stimulation, multiple small vacuoles within the hyphal cytoplasm merge into larger compartments. TEM images showing vacuolar distribution before (**G**) and after a 45 °C, 10-min thermal stimulation (**H**) (scale bar, 2 μm). **I.** 1D ^13^CP spectra (left) and ^13^C spectra refocused INEPT (right) solid state NMR results when heating the living fungal from 23 °C to 55 °C and back to room temperature. **J.** Response of a single-channel Mycoelectronics thermoreceptor under 1 minute of temperature stimulation from 25 °C to 45 °C. **K.** Response of a single-channel Mycoelectronics under 5 minutes of temperature stimulation from 25 °C to 45 °C. **L.** Thermal response of the Mycoelectronics under various temperatures showing the ability to distinguish 2 °C temperature differences. **M.** Quantitative analysis of the relationship between thermally induced voltage changes and temperature for the Mycoelectronics. **N.** Comparative analysis of four independently prepared Mycoelectronics, demonstrating consistent voltage responses under 25 °C, 35 °C, and 45 °C thermal stimulation. **O, P.** Spatial resolution capabilities of the single-channel Mycoelectronics under localized heat source positioning. **Q.** Long-term stability of a Mycoelectronics under cyclic stimulation at (room temperature for 10 min, then 35 °C for 1 min).

Chronopotentiometry measurements show that the live mycelium exhibits repeatable thermoresponsive behavior. In contrast, control experiments show that dead mycelium, obtained by heat-killing and subsequently sealed to maintain controlled humidity, or bare culture medium, do not exhibit such changes, as they lack the active cellular processes necessary for signal generation (Fig. 3E), indicating that biological viability is essential for the thermal response. To understand the sensing mechanism underlying the heat-induced changes in the electrical properties of the mycelial network, we employed a set of optical microscopies, electron microscopy, and solid-state NMR (ssNMR) techniques to characterize the structural and compositional dynamics during thermal stimulation. Transmission electron microscopy (TEM), along with confocal fluorescence microscopy and time-lapse imaging, revealed rapid vacuolar remodeling and fusion in response to thermal stimulation. Specifically, thermal elevation resulted in a significant increase in vacuole size, indicating intracellular osmotic and metabolic remodeling (Fig. 3F). The resulting compartmentalization of the conductive cytoplasm by lipid-bound vacuoles reduces the effective volume of ionic conduction. Compared with the no-thermal-stimulation condition (Fig. 3G), 10 min of thermal stimulation triggered vacuolar fusion (Fig. 3H), producing a single large compartment that disrupts continuous ion pathways and restricts ion flow. Systematic TEM characterization across varying thermal stimulation conditions, with subsequent resting at ambient temperature, reveals the rapid and highly reversible nature of vacuolar fusion and remodeling (fig. S12). Previous studies suggest that fungal thermal responses are partially governed by thermal stimulation proteins (HSPs)(*76*, *77*) such as HSP 90 and HSP 101, as well as upstream transcription factors like HsfA(*78*, *79*), which rapidly regulate gene expression in response to sudden temperature changes. This transcriptional response can occur within minutes, aligning with our observation of abrupt shifts in electrical resistance upon thermal stimulation. In addition to transcriptional regulation, fungi accumulate thermo-protective osmolytes such as trehalose and glycerol to retain intracellular water and mitigate heat stress. Internal resources are also reorganized; lipid droplets proliferate to modulate membrane lipid composition and sequester stress-induced byproducts(*80*). Consistent with this, we observed a rapid increase in lipid droplet size within minutes under real time confocal microscope, as shown in fig. S13 and Movie 6.

Supporting evidence from ssNMR further reveals heat-induced alterations in both lipid and protein content within the cell wall, followed by partial recovery after the stress is removed. The one-dimensional ^13^C cross-polarization spectra detecting rigid molecules showed significant intensity changes corresponding to lipids and proteins (Fig. 3I, left), reflecting structural remodeling in the load-bearing regions of the fungal network(*81*, *82*). In contrast, the refocused INEPT experiment spectra detecting highly mobile molecules (Fig. 3I, right) did not show significant changes in either the carbohydrate region (Fig. 3I, right) nor the lipid and protein part (fig. S14). These findings are consistent with our reported structural reorganization of fungal cell walls induced by thermal stress, which includes the loss of cell wall crosslinking, structural loosening, and altered dielectric behavior in fungal tissues. Together, the electrical and spectroscopic data indicates that heat-induced structural reorganization—particularly vacuole fusion and remodeling within conductive cytoplasmic domains—contributes to a gradual and measurable increase in mycelial electrical resistance.

Building on these mechanistic understandings and insights, we next characterize the sensing performance of the living system. Stepwise heat stimuli were given from 23 °C to 45 °C, and the Mycoelectronics thermoreceptor revealed a clear voltage response corresponding to each temperature increment (Fig. 3J). Under prolonged heating conditions of 5 minutes at each temperature level, the system exhibited adaptation responses due to the living nature of mycelium (Fig. 3K). Under different heat stimuli, the voltage-time data shows a quick response (less than 2 seconds) and the discern temperature can be identified as approximately 2 °C. (Fig. 3L). Fig. S15 and Movie 7 demonstrate rapid changes in mycelial fluorescence intensity in response to environmental temperature variations. Importantly, there is a relative linear relationship between voltage output and heat stimulus temperature (Fig. 3M). The increasing electrode distance compromises the thermal sensing capabilities, resulting in defined effective and ineffective temperature sensing regions as shown in fig. S16. Four independent samples show relatively minor voltage variations with small fluctuations (Fig. 3N), indicating the fungi bioelectronic thermoreceptor system’s potential for precise temperature quantification. The single-channel Mycoelectronics thermoreceptor can be used for spatial temperature discrimination, distinguishing temperature differentials across a 3×3 sensing array, with notably higher thermal response detected at position 21 compared to surrounding regions (Fig. 3O, P). Remarkably, the fungal sensor maintained stable performance over 300 cycles across 55 hours (Fig. 3Q), underscoring the potential of Mycoelectronics for long-term and reliable thermal response as a biohybrid sensation system. Our extended studies further show that the living Mycoelectronics deliver stable performance for at least 8 consecutive days (fig. S17). Moreover, when stored under controlled humidity, they retain sensing capability for at least two months without replacing the growth medium, requiring only minimal recalibration before use (fig. S18).

The mycelial network’s ability to grow, assemble, heal itself, and navigate through physical constraints enables sensing capabilities in architecturally complex or hard-to-reach locations where traditional electronic systems would be impractical or even impossible to implement. To characterize these properties, we first validated the self-healing capacity of the Mycoelectronics thermoreceptor through mechanical damage (Fig. 4A and fig. S19). The experiment began with the fungal inoculation on a gelatin-based medium, followed by the establishment of an active response network between electrodes. Within three days, the mature mycelium network was deliberately damaged with a razor blade. Following the damage, the fungal system demonstrated rapid self-healing by regenerating new mycelial connections across the damaged area (Movie 8). Even after disrupting a 1 cm-wide aerial mycelium, the ion-electronic pathway was restored in as little as two hours (fig. S20). By day 7, the regenerated network maintained its thermal responsiveness with minimal attenuation compared to the original baseline (Fig. 4B).

**Fig. 4.**
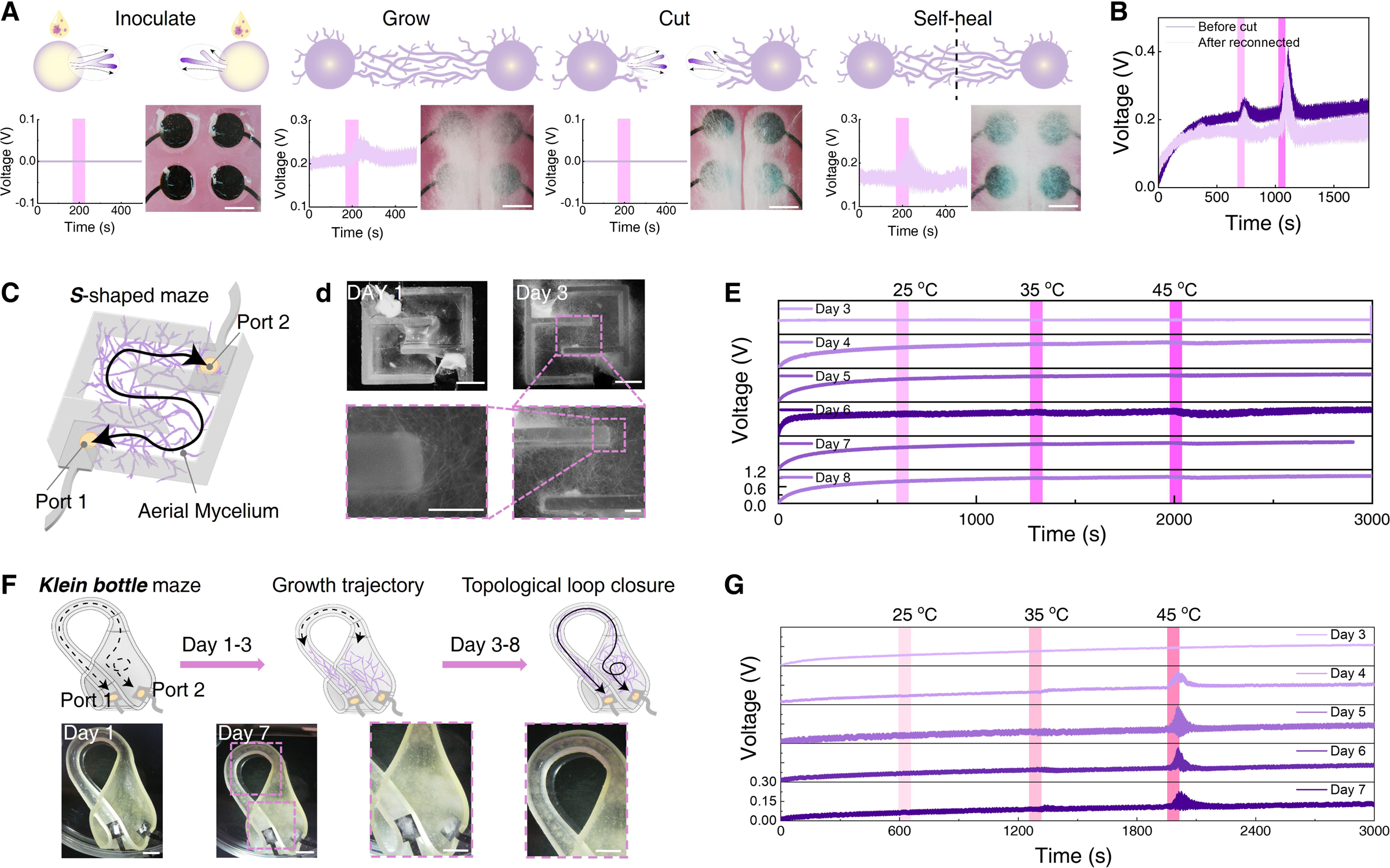
Self-healing, adaptive growth and assembly of the Mycoelectronics. **A**, Schematic and experimental demonstration of the self-healing capability of Mycoelectronics to rebuild the connection and sensing functions. The top row shows schematic illustrations; the bottom row shows corresponding optical microscope images and their electrical signals (scale bar, 5 mm). **B,** The voltage response of the fungal network thermal response before mechanical damage and after regeneration. **C,** Schematic of the adaptive growth and assembly of the fungal network connection in an S-shaped PDMS maze model. **D,** Optical images of connected fungal growth in S shape maze (scale bars, top 1 cm; bottom: 1 mm). **E,** Daily temperature response of the Mycoelectronics in the S-shaped maze during the mycelium growth and navigation. **F,** Schematic and photographs of the adaptive growth and assembly of the fungal network along a Klein bottle maze, achieving connection after 7 days (scale bar, 1 cm). **G,** Daily temperature response of the growing Mycoelectronics results in the Klein bottle maze during the mycelium growth and navigation.

The fungal bioelectronic thermoreceptor system exemplifies the synergy of biological intelligence and electronic functionality, as the mycelial network adapts its architecture through tip-guided extension and context-dependent branching —a crucial feature for future intelligent and autonomous circuit design and manufacturing. To illustrate this capability, two fungal clusters were inoculated at the ends of an *S*-shaped maze (Port 1 and 2) (Fig. 4C). Over three days, the network autonomously navigated the maze and assembled a conductive sensing pathway (Fig. 4D), with increasing thermal response observed over time (Fig. 4E). In a more topologically complex half, *Klein* bottle maze (Fig. 4F), the mycelium successfully grew through the only available channel and formed a functional connection between Port 1 and Port 2 within one week, as confirmed by subsequent thermal signal generation (Fig. 4G). These findings underscore the potential of living fungal electronics to enable self-assembling, self-healing, and self-routing sensory networks for intelligent infrastructure and distributed biosensing in physically constrained environments.

We then explore the potential applications of the fungal bioelectronic thermoreceptor, including environmental sensing, artificial nerve, and robotics. The comprehensive signal processing workflow for living Mycoelectronics, enabling artificial sensation and response, is illustrated in Fig. 5A. The process can be divided into three main stages: signal detection, signal processing, and signal output for various applications. Raw voltage signals from the sensor array undergo chronopotentiometry and filtering before entering three key processing modules: amplitude detection, spike detection and voltage comparison, which finally enable various applications. Real-time processing was achieved using PalmSens4 hardware for voltage readings and an Arduino board for signal comparison against preset thresholds, triggering appropriate circuit controls when critical values were reached. In our multi-channel sensing platform, the mycelial network functions as a distributed temperature sensor array, transmitting various voltage values through a flexible flat cable to downstream devices (Fig. 5B). Upon localized heating, infrared thermal imaging revealed distinct spatial patterns of temperature distribution, with voltage amplitudes correlating with local thermal gradients (Fig. 5C). The lower-left panel of Fig. 5c shows a 3D bioprinted 16-channel Mycoelectronics array. To generate high-resolution thermal maps from discrete sensor data, cubic spline interpolation was applied to signals from the eight sensing nodes, producing continuous temperature-distribution profiles (fig. S21). The mechanical flexibility and spatial thermal mapping capacity were demonstrated to meet omni-directional temperature monitoring demands on a 3D surface (Fig. 5D). A distributed 8-channel Mycoelectronics on a sphere demonstrates capability of identifying the location and intensity of the heat source, as shown in Fig. 5E, F and fig. S22. We envision such sensors could be used for wildfire monitoring. This self-organized growth of the fungal network significantly increases the effective sensing surface area. It thus reduces the need for intricate circuit designs.

**Fig. 5.**
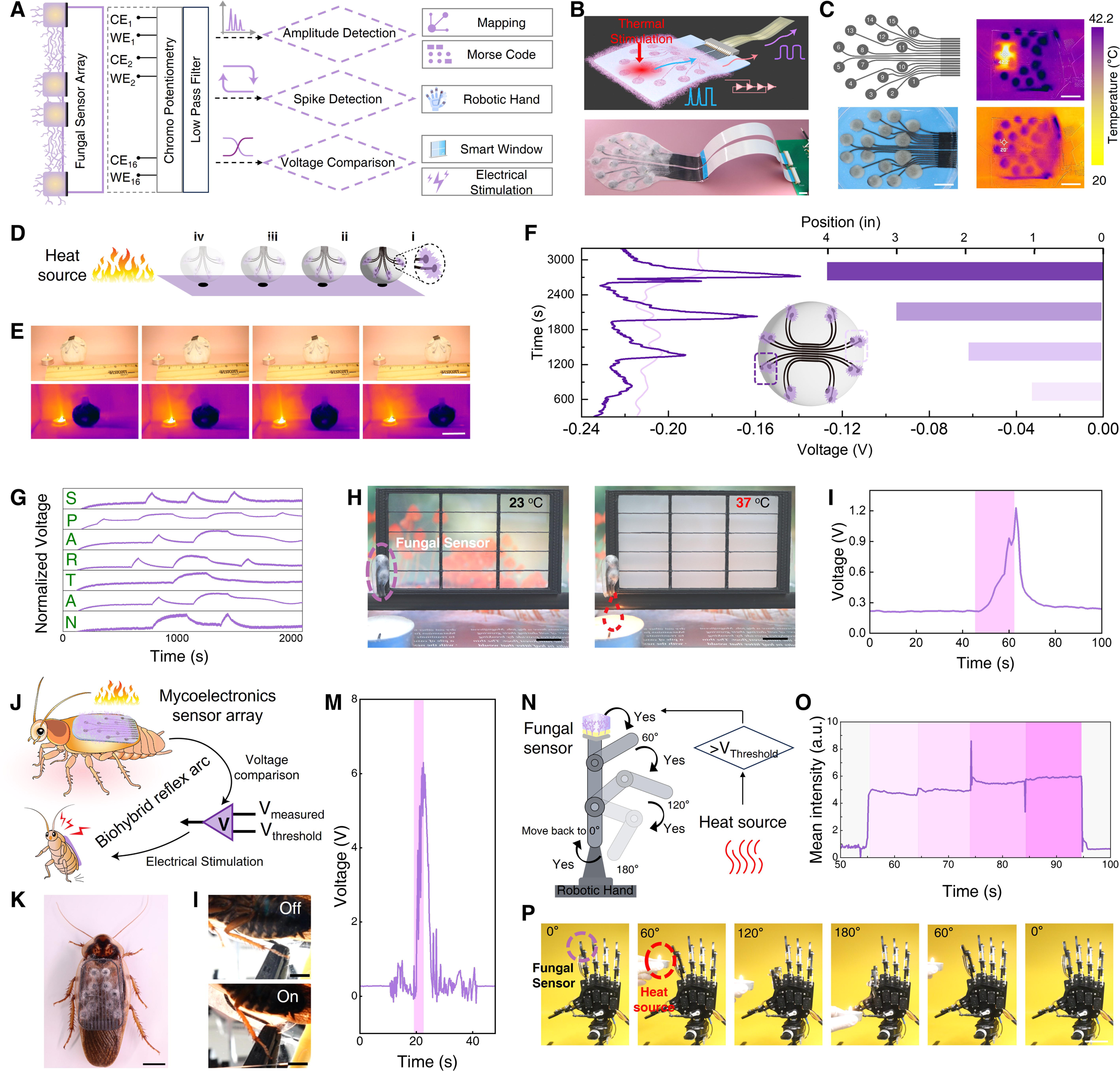
Applications of Mycoelectronics thermoreceptor in environmental sensing, and artificial sensation for animals and robots. **A**, Signal processing architecture showing three pathways of the Mycoelectronics thermoreceptor for amplitude detection, spike detection, and voltage comparison, enabling diverse output functionalities. **B,** Schematics and photography of the multi-channel Mycoelectronics integrated with signal readout capabilities (scale bar, 1 cm). **C,** Design and thermal characterization of the multi-channel Mycoelectronics for thermal mapping. Left: schematic and real image of the 16-channel electrode layout. Right: thermal imaging demonstrating spatial temperature distribution (20 °C-42 °C) (scale bar, 2 cm). **D,** Schematic of 3D thermal mapping experiment setup. **E,** Optical (upper) and infrared (bottom) images portfolio of 3D thermal mapping using the multi-channel fungi Mycoelectronics to identify the location and intensity of the heat source (scale bar, 1 cm). **F**, Two representative temperature responses (closest and farthest to the heat source) results of the 3D Mycoelectronics temperature sensor array. **G,** Temperature-based Morse code transmission by Mycoelectronics. **H,** Optical images showing the Mycoelectronics to control a PDLC window with and without thermal stimulation (scale bar, 1 cm) and **I,** Corresponding Mycoelectronics voltage response. **J,** Schematic showing the process to engineer Mycoelectronics as a thermoreceptor for a cockroach to build the thermal sensation and muscle actuation functions: electrode fabrication, fungal printing, and responsive bioelectronic stimulation. **K,** Image showing a Dubia cockroach with a Mycoelectronics sensor array on its back for thermal sensation (scale bar, 1 cm). **L,** Photographs showing the cockroach with thermally induced leg motion mediated by the Mycoelectronics thermoreceptor (scale bar, 5 mm). **M.** Voltage response of the Mycoelectronics on cockroach’s back, pink background marks the duration when the heat source was positioned near the sensor. **N, O.** Schematic and biomechanical control of robotic hand with integrated Mycoelectronics thermal sensors (scale bar, 5 cm). **P,** Sequential images demonstrate thermotactic behavior with the microelectronic thermal sensation (scale bar, 5 cm).

Ubiquitously distributed in nature, fungi possess innate harmlessness and camouflage. Mycoelectronics thermal fluctuations could be translated into a voltage signal response duration and further reversely encoded information using Morse code. A message ‘SPARTAN’ was transmitted by thermal-to-electrical signal encryption (Fig. 5G). We then installed our Mycoelectronics on the outer surface of a building, the living thermoreceptors provided binary output to control smart windows in response to environmental temperature changes, providing potential for smart energy-efficient building management (Fig. 5H, I and Movie 9).

We engineered the living Mycoelectronics system as a wearable electronic skin, using the cockroach as a model animal, with artificial afferent pathways to emulate the function of biological thermoreceptors and sensory nerves. A multi-channel Mycoelectronics array was designed and fabricated into a wearable device that mimics the distributed tactile receptors of natural skin. To create a Mycoelectronics biohybrid reflex arc, this integrated system combines fungal–electrode interfaces, mycelium-based signal generation, and downstream electronic stimulation with wire electrodes. Upon thermal activation, the system digitizes bioelectrical signals and triggers stimulation of the cockroach leg (Fig. 5J, K). Once the heat-induced signal exceeded the preset voltage threshold of the Mycoelectronics, electrical stimulation was delivered to the cockroach leg, resulting in contraction. Upon cooling, the voltage dropped below the threshold, and stimulation was terminated, allowing leg relaxation (Fig. 5L, M and Movie 10). These findings demonstrate the potential of Mycoelectronics to serve as biohybrid skins for environmental sensing, adaptive response, and escape behaviors. Extending this concept, we integrated a fungal thermoreceptor onto the fingertips of a robotic arm. Upon exposure to heat from a candle flame, the mycelium network generated real-time signals to command the robotic arm to keep moving, enabling autonomous, heat-responsive behavior governed by living fungal intelligence (Fig. 5N, O and Movie 11). Together, these results establish a living Mycoelectronics platform that bridges thermal sensation and reflexive actuation, offering a new class of biohybrid systems for neurorobotics and neuroprosthetic applications.

## Discussion

The quest to replicate biological sensation has long challenged engineers and scientists, and the biohybrid strategy offers a compelling alternative. We introduced fungal *Mycoelectronics* as a living biohybrid platform that integrates fungal mycelium with stretchable graphene-based electrode array to mimic biological thermal sensation. The artificial sensation arises from a newly discovered mechanism: heat-induced vacuole remodeling within living fungal cells, which modulates rapid and reversible changes in their electrical conductivity. Unlike conventional biohybrid systems that rely on mammalian cells or bacteria, which require stringent culture conditions and are often limited in scalability, this system achieves seamless biological-electronic coupling through a scalable bioprinting process and the subsequent biological growth of the mycelium networks. This artificial sensation system leverages fungi’s unique biological properties: they are environmentally responsive, naturally electrically conductive, self-healing, and self-adaptive through rapid proliferation. Comprehensive characterizations and proof-of-concept demonstrations— covering single-channel and multi-channel sensing, self-healing ability, topography solving, and closed-loop control—reveal the unique robustness and versatility of the sensor.

Our results highlight several key innovations. First, we demonstrate that living fungal networks can act as real-time thermoreceptors, with thermal responsiveness emerging from intrinsic biological processes such as vacuolar remodeling. The fact that such cellular-level events can manifest as measurable electrical changes bridges the gap between molecular biology and electronic function. Second, we establish a scalable, reproducible biofabrication strategy for living bioelectronic Mycoelectronics. Graphene based electrodes patterned on soft substrates are directly integrated with a nutrient-rich fungal bioink via additive bioprinting. This approach preserves biological viability, allowing motile, self-organizing mycelial networks to form functional sensor architectures capable of autonomously growing into hard-to-reach environments. Third, we demonstrate a universal signal processing flow that converts fungal thermal responses into discrete electronic outputs. This enables not only practical sensing applications—such as thermal encryption and smart building regulation—but also the development of cross-kingdom communication systems. Notably, we establish a functional reflex arc by interfacing fungal signals with cockroach neuromuscular systems and further extend this to robotic actuation, enabling autonomous, heat-triggered movement.

Finally, the potential of fungal bioelectronics extends beyond thermal sensing. The inherent adaptability, resilience, and inconspicuous natural morphology of fungi make them ideal candidates for next-generation, environmentally integrated electronics. Future studies may expand their use to detect diverse stimuli and environmental cues(*56*, *58*, *61–63*), and explore their role in ecological monitoring, wearable systems, and even implantable sensing systems, given the abundance of fungi and their significant impact on human health(*59*, *60*, *83*). In summary, our work lays the foundation for standardized fungal-electronic systems and reveals a path toward multifunctional, sustainable, and adaptive living sensors. Fungal Mycoelectronics offer not only technological utility but also a fundamentally new perspective on integrating natural systems with engineered ones.

## Materials and Methods

### Materials

The following chemicals were purchased and used as received: malt extract (Carolina), agar (Sigma-Aldrich), yeast (Sigma-Aldrich), Gelatin from porcine skin (Sigma-Aldrich), Rhodamine (Sigma-Aldrich), Fluorescein sodium salt (Sigma-Aldrich).

### Preparation of growth media and cell culture

Two types of growth media were prepared as feeding substrates for the fungus: malt extract agar (MEA) and Gelatin based ME. MEA medium was prepared by adding 20 g/L (2%) malt extract, 2 g/L (0.2%) yeast and 20 g/L (2%) agar. To obtain MEA plates, the medium was sterilized at 121 °C for 45 min in an autoclave (2840 EL, Tuttnauer) and, subsequently, poured into cell culture dishes and left to solidify. The Gelatin base MEA substrate was prepared by mixing 1:6 gelatin to water at 60 °C for 1 hour.

The fungi were subcultured by placing a small slice of a MEA plate fully covered with mycelia onto a fresh growth plate (2% malt extract) every 3–5 weeks. The cultures grew in the dark at 23 °C and 30% relative humidity.

### Electrode fabrication

The electrode was made by transfer-printing laser-induced graphene (LIG) electrodes onto Styrene Ethylene Butylene Styrene (SEBS) (Texin RxT85A) elastomer substrates. Specifically, the LIG electrodes were engraved on a polyimide film using a 30 W CO_2_ laser cutter (power, 15%; speed, 25%; VLS2.30 universal laser systems). Next, SEBS solution (0.2 g/ml in cyclohexanone) was applied on LIG electrode and dried at room temperature. After drying, the SEBS layer containing the LIG electrodes was carefully peeled off from the polyimide substrate. Finally, to encapsulate and protect the electrode leads, a solution of SEBS in acetone (0.05 mg/ml) was applied orthogonally to the electrode pattern.

### Bioprint ink preparation

The fungal bioink was prepared by dissolving malt extract (12 g) and agar (9 g) in distilled water (200 mL). The mixture was autoclaved at 121°C for 20 minutes and allowed to cool to room temperature. To achieve optimal printing viscosity, the solidified medium was mechanically homogenized using a laboratory blender. The processed medium was then transferred to sterile Petri dishes and inoculated with *Benniella eriona (MycoBank No: MB#833779)*. After one week of cultivation at 23°C, the aerial hyphae were carefully removed, and the remaining substrate containing established mycelial networks was harvested. The resulting fungal-medium composite was gently transferred to sterile printing cartridges. Three-dimensional printing was performed using Brinter® ONE (Brinter Ltd., Finland) with predetermined geometric patterns. Printing parameters were optimized with extrusion pressure of 500 kPa and printing speed of 8 mm/s to ensure consistent material deposition and maintain mycelial viability.

### Gap Bridging Setup

Each fungal sample (a cylindrical block with a 1 cm² cross-sectional area and around 5 mm height) was transferred to an empty sterile Petri dish, placing two blocks at defined distances apart (1 cm, 2 cm, 4 cm, 6cm, 8 cm and 10 cm). Plates were incubated at 23°C and 30% RH. Successful bridging was defined as visible aerial mycelium fully connecting the two fungal blocks.

### Single channel Mycoelectronics fabrication

1 cm × 1 cm square LIG electrode was created using laser induction techniques. Conductive traces are made of LIG also, measuring 5 cm in length and 2 mm in width. Then using SEBS to form a stretchable LIG electrode. Two such LIG electrodes were positioned with a standard spacing of 1 cm between them. On each electrode, a medium block (1 cm x 1 cm in area and 5 mm thick) containing fungi was placed. The parafilm was used to maintain a controlled environment and prevent contamination. After three days of incubation, a single-channel sensor was ready. A 1 nA current was applied using PalmSens and equilibrate for 5 min before introducing any thermal stimulus.

### Multichannel Mycoelectronics fabrication

Multichannel electrode array was designed using Adobe illustrator, then the LIG method to fabricate. After transferring on stretchable SEBS substrate, a gelatin-based MEA solution was drop-casting onto the LIG electrodes at the same area. The entire assembly was sealed with parafilm and incubated for three days. After incubation, the agar-based fungi were removed, and the fungi were allowed additional time to naturally form connections between the multichannel electrodes.

### Electrochemical Impedance Spectroscopy (EIS) test

A two-electrode system (Palmsens) was used by placing the fungus between two LIG electrodes. Choose an AC signal of 5mV with frequencies from 1Hz to 100000Hz.

#### Conductivity test

EIS testing was conducted to record impedance. Identify the resistance value (R) when the imaginary part of the impedance (EIS imaginary component) is zero. Measure the length (L), width (W), and height (H) of the sample. Calculate the ionic conductivity of the gel and the gel-mycelium composite material using the following formula:

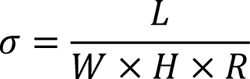

### Mechanical test

Mechanism Test was performed using a CellScale (UniVert) mechanical testing system in tension mode. Force and displacement were recorded continuously until sample failure or reaching the set limit. Data were converted to stress–strain format assuming σ = F/A and ε = ΔL/L_0_.

Loop tack experiment was conducted with Instron 5565 Tensile tester equipped with Instron Series 2712 Pneumatic Grips. Strips approximately 8 × 40 mm in size were cut for loop tack testing, and a drop loop was formed by taping together and clamping 10 mm of each end of the strip in the upper grip of the tester. The upper grip was moved downwards at a rate of 1 mm/s until an area of 8 × 8 mm of the graphene substrate was covered. Once this area was covered, the upper grip was moved upwards at a rate of 5 mm/s and the maximum force per meter required to remove the specimen loop from the substrate was recorded as the loop tack.

### Rheology test

The rheological behavior of bioink and hydrogel was measured by Discovery HR-2 rheometer (TA Instruments, USA). All measurements were performed at room temperature, and data were processed using TRIOS software (TA Instruments, USA). Apparent viscosity was recorded over a shear rate range of 0.1 to 100 s⁻¹ to assess shear-thinning behavior under steady shear. Three-interval thixotropy test (3ITT) for bioink and hydrogel behavior were assessed by alternating oscillatory (1% strain, 10 rad/s) and steady shear at 1 s⁻¹ in repeated cycles. Each cycle included 20 seconds of oscillatory measurement to establish the baseline moduli, followed by 30 seconds of steady shear to simulate extrusion. Oscillatory strain sweep tests were performed at a constant angular frequency of 10 rad/s with strain amplitudes ranging from 0.1% to 100% to determine the storage modulus (G′), the loss modulus (G″).

### Scanning Electron Microscope (SEM) image

To prepare the biological samples for SEM, a critical point drying (CPD) process was performed. The biological samples were fixed in 4% glutaraldehyde in 0.1 M sodium phosphate buffer for 1 hour. After fixation, they were rinsed buffer for 30 minutes. The samples were then dehydrated through a graded ethanol series (25%, 50%, 75%, 95%, and three changes of 100% ethanol, 10 minutes per step). Once in 100% ethanol, the samples were transferred to a critical point dryer, where they were dried for over 2 hours. After drying, the samples were mounted and coated for further analysis.

A scanning electron microscope (JEOL Ltd., Tokyo, Japan) was used to image the morphology of multiple samples. Gold coating was applied before imaging with the SEM at 10kV.

### Transmittance Electron Microscope (TEM) image

Specimens were chemically fixed in 2.5% glutaraldehyde in 0.1 M phosphate buffer (pH 7.4) for 2 h, rinsed thoroughly, and post-fixed in 1% osmium tetroxide for 1 h at room temperature. Following dehydration through a graded ethanol series and infiltration with epoxy resin, samples were embedded and polymerized. Ultrathin sections (70 nm) were sectioned using an ultramicrotome, mounted on copper grids, and contrasted with uranyl acetate and lead citrate. Transmission electron microscopy was performed at 80 kV (JEOL/FEI system) to capture both longitudinal and transverse views of the fungal cellular architecture.

### Confocal microscopy image

The fungal mycelium was stained with [Biotium DIBAC_4_(3)], 2µM. After staining, the samples were washed with PBS to remove any excess dye. These were then imaged with CLSM (TCS SP8 X, Leica, USA)

### Solid state NMR test

Solid-state NMR experimental procedures: The 1D high-resolution solid-state NMR experiments were conducted on a Brucker Advance 800 MHz (18.8 Tesla) spectrometer using 3.2 mm rotor under 15 kHz MAS at 298 K. The ^13^C chemical shifts were externally referred to adamantane CH_2_ signal at 38.48 ppm on the TMS scale. The ^15^N chemical shifts were referred externally through 110.0 ppm of Phe peak in the MLF model peptide. The magic angle was calibrated using KBr. Typical ^1^H radiofrequency field strengths 50-83 kHz and 50-62.5 kHz for ^13^C. The initial magnetization for the experiment was created using ^1^H-^13^C cross-polarization that preferentially detects rigid molecules. Typically, 1 ms Hartmann-Hahn contact was used for the CP. The J-coupling-based ^1^H-^13^C refocused Insensitive Nuclei Enhancement by Polarization Transfer (INEPT) experiment(*84*) targeted the most mobile molecules by using J-coupling to transfer magnetization between bonded ^1^H and ^13^C nuclei. This approach efficiently detects highly mobile molecules with long transverse relaxation times during four delays of 1/4J_CH_, 1/4J_CH_, 1/6J_CH_, and 1/6J_CH_ in the pulse sequence, where J_CH_ represents the carbon-hydrogen J-coupling constant and was set to 140 Hz. Lastly, 1D ^13^C direct polarization (DP) experiment with a short recycle delay of 2 s was employed to preferentially detect mobile molecules with fast ^13^C-T_1_ relaxation.

### Maze channel fabrication

The Fusion 360 designed maze model was imported to Form3 3D printer (Formlabs Inc.). High Temp Resin was used for the maze model. Cover this model with a mixture of Polydimethylsiloxane (PDMS) base and curing agent in a 10:1 ratio, and then cure it at 60°C for 2 hours to form a solid PDMS mold. Carefully demold the cured PDMS to reveal the maze channels. Place two LIG electrodes at the entrance and exit of the maze. On each electrode, place a drop of gelatin-based medium, and then introduce the mycelium on top. Over the following days, capture daily photographs of the maze and measure the electrical signals between the two electrodes(*76*).

### PDLC window control

The setup consisted of a Polymer Dispersed Liquid (PDLC) window panel connected to an external power supply via a relay, which was controlled by an Arduino Mega board. The fungal sensor was placed near the window and connected to a PalmSens electrochemical interface for voltage measurements. Real-time data was acquired using Tesseract OCR and processed by a custom Python script. The script continuously compared the sensor’s voltage output to a preset threshold corresponding to a specific temperature. When heat was applied and the voltage exceeded the threshold, the Arduino triggered the relay to cut power to the PDLC window, turning it opaque. Conversely, as the temperature decreased and the voltage fell below the threshold, power was restored, returning the window to its transparent state.

### Robotic arm reflex arc

A pair of bioelectrodes is employed to capture the physiological or environmental signals, and this single-channel sensing device is attached to the outer side of the robotic arm’s index finger, ensuring optimal contact and signal acquisition. The entire system is interfaced with an Arduino Mega 2560 board, which serves as the central control unit for signal processing and provide PWM wave as the actuator command execution, the power supply including the positive and negative signals are independently provide by external DC power source and isolated from the control board.

When the sensor detects a rising voltage which indicates the thermal stimuli, the system interprets this as a trigger for motion initiation. As the input voltage continues to increase, the Arduino sequentially commands the servo motors of the robotic arm to actuate the finger. The movement progresses through distinct angular positions using a open-loop control: first from the rest position at 60°, then incrementally to 120°, and further to 180°, representing a fully extended state. This staged actuation is designed to mimic an intelligent behavioral response, wherein the robotic arm is actively retracting or repositioning itself in response to an external heat source. After completing the motion, the system resets the arm position back to 0°, preparing it for the next cycle of sensing and response.

### Electrode implantation

Our method of steering the insect’s locomotion was based on existing procedures(*85*, *86*). Prior to surgery, insects were anesthetized by cooling with ice packs in a Styrofoam cooler for around 30 minutes. The insect needle was inserted into the leg until feeling noticeable resistance. The ground electrode was inserted at the midline through the second abdominal segment via a small dorsal puncture made with an insect pin. The stimulation electrode was inserted into the base of the cockroach’s leg. The test subject was allowed to recover under a warming lamp for about 30-60 min.

### Stimulation of cockroach leg

A square wave signal was generated and formed a symmetric on/off waveform suitable for neuromuscular stimulation with precisely optimized parameter, which are amplitude of 2.5 V, frequency of 50 Hz, and 50% duty cycle. This signal was first produced using the internal timer modules of the Arduino Mega 2560, eliminating the need for external waveform generators. To ensure the signal reached the required amplitude for effective stimulation of the cockroach leg muscles, the output from the Arduino was passed through an operational amplifier circuit based on OPA451 structure. This amplification stage adjusted the digital output into a finely controlled signal with an amplitude precisely regulated to 2.5 V peak, maintaining signal integrity and minimizing distortion during transmission. This stimulation was directed to the leg muscles of the cockroach. A fungal temperature sensor was attached to the cockroach’s back using adhesive. PalmSens was used to conduct a constant current test at 1nA. The real-time voltage from PalmSens was captured using Tesseract OCR technology and transmitted to an Arduino. The feedback part of this system included a voltage-sensing module connected to one of the analog input channels on the Arduino Mega. This external sensor continuously monitored the voltage level output from the single-channel detection device. A predefined threshold voltage was set in the Arduino firmware; when the input signal from the sensor exceeded this threshold, it triggered the output of the square wave signal to initiate electrical stimulation of the cockroach leg muscles. To prevent false triggering due to signal fluctuations or noise, a hysteresis control scheme was implemented by defining a 0.2 V buffer window around the threshold.

## Supporting information

Supplementary Video 1 Fungal aerial mycelium growing toward and interacting with LIG electrodes

Supplementary Video 2 Loop tack test of gelatin based medium

Supplementary Video 3 Fracture dynamic of a freestanding fungal mycelium network

Supplementary Video 4 3D bioprinting of bioink patterns on a LIG electrode array

Supplementary Video 5 Synchronized optical, infrared, and electrical data acquisition of mycoelectronics thermoreceptor

Supplementary Video 6 Real-Time observation of vacuolar fusion in hyphae

Supplementary Video 7 Voltage sensitive fluorescence response of mycelium to thermal stimulation

Supplementary Video 8 Rapid aerial mycelium network formation

Supplementary Video 9 Temperature-triggered smart window switch using a mycoelectronics thermoreceptor

Supplementary Video 10 Mycoelectronics sensor activates cockroach leg motion

Supplementary Video 11 Mycoelectronics sensor activates robotic hand motion

## Acknowledgments

**Funding:** J.L. acknowledges support from the National Science Foundation under Award Nos. ECCS 2339495, ECCS-2334134, ECCS-2216131, EFMA-2318057, and CMMI 2323917. Solid-state NMR analysis was supported by NSF MCB-2517270 to T.W.

## Author contributions

Conceptualization: JL, YC, VT. Methodology: YC, CR, VT, HY, LX, SK. Investigation: JL, TW, GB. Visualization: YC, HY, SK.

Supervision: JL, TW, GB.

Data collection: DF, SK, IS, KZ,

Writing—original draft: YC.

Writing—review & editing: YC, CR, HY, KS, LX, LH, KN, BW, TW, GB, JL.

## Data and materials availability

All data are available in the main text or the supplementary materials.

## Supplementary Materials

Figures. S1 to S22

Movies S1 to S11

