## Supplementary Information for "Mycoelectronics: Bioprinted Living Fungal Bioelectronics for Artificial Sensation"

Yulu Cai *et al.*

**The PDF file includes:**

Figs. S1 to S22

Legends for movies S1 and S11

**Other Supplementary Material for this manuscript includes the following:**

Movies S1 and 11

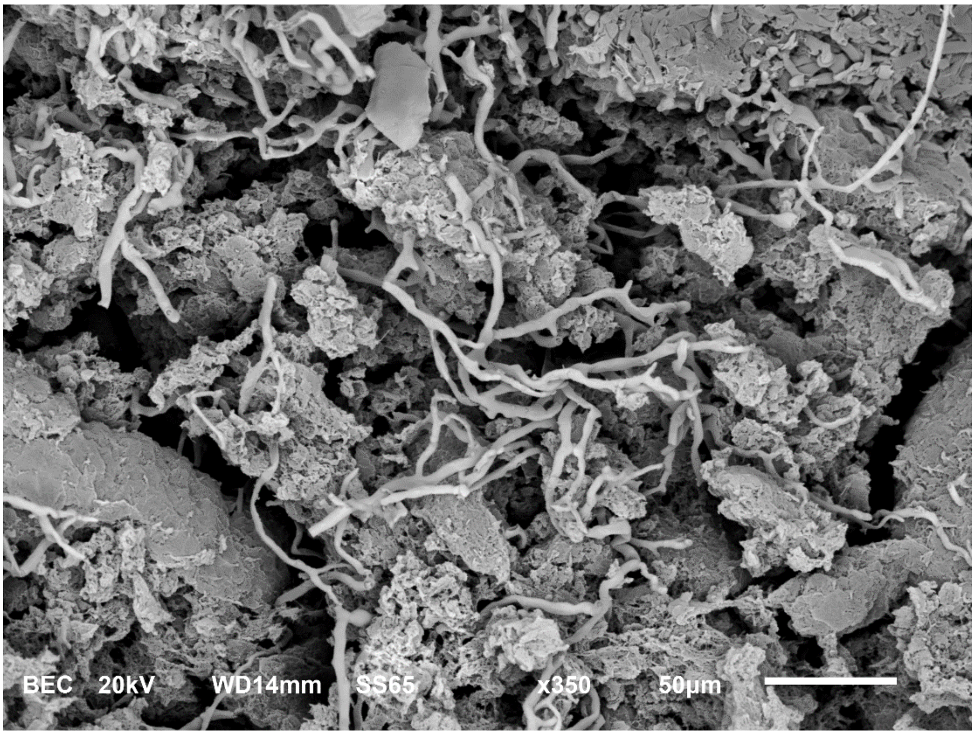


**Fig. S1. Backscattered Electron Composition image of Fungi bioink.** Backscattered Electron Composition characterization of the bioink interface demonstrates a clear layered architecture, where the dense *Benniella erionia* fungal mycelium network forms intimate contact with the culture medium layer.

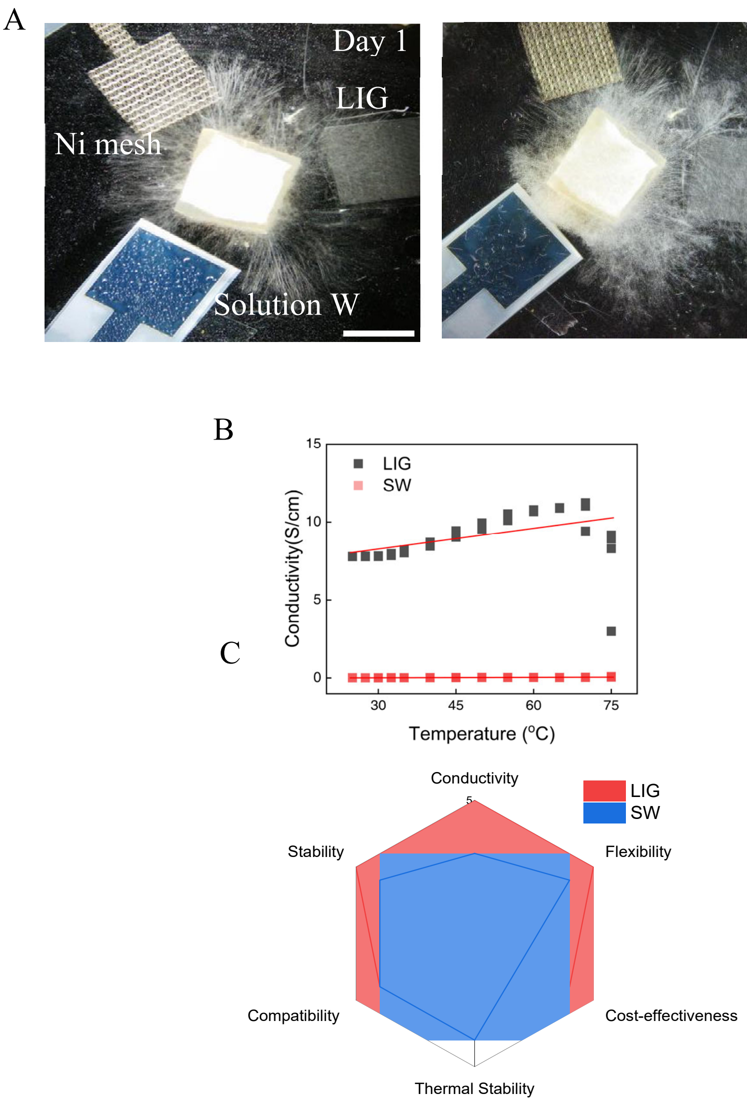


**Fig. S2. Electrode selection.** (**A**) Electrode selection for fungal bioelectronics, comparing LIG and SW (PEDOT:PSS based material), and Nickel (Ni) electrodes in reactivity with *Benniella erionia*. The top row presents time-lapse images over two days, demonstrating superior fungal growth on LIG electrodes compared to SW and Ni (scale bar, 1 cm). (**B**) Conductivity versus temperature, with LIG maintaining higher conductivity across temperatures from 30°C to 75 °C, while SW exhibits negligible conductivity. (**C**) A radar chart on the bottom right compares six key properties: stability, conductivity, flexibility, thermal stability, compatibility with medium, and cost-effectiveness. LIG outperforms SW in most categories, particularly in conductivity, stability, and medium compatibility.

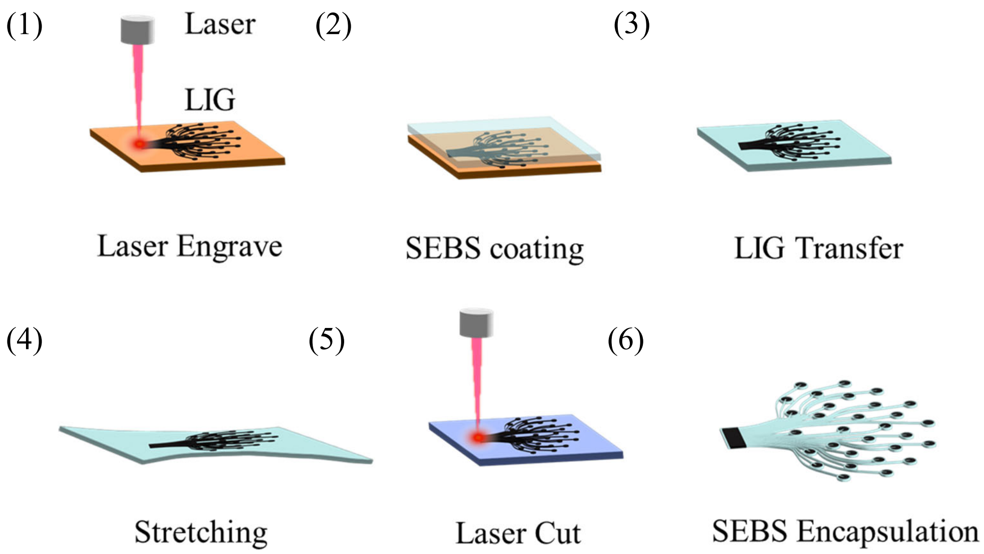


**Fig. S3. Fabrication process for self-supporting, stretchable electrodes using LIG.**

The method comprises six key steps: (1) Laser Engrave, where a laser creates a LIG pattern on a substrate; (2) SEBS Coating, involving the application of a Styrene-Ethylene-Butylene-Styrene layer over the LIG; (3) LIG Transfer, where the coated LIG is peeled off and flipped onto a new substrate; (4) Stretching of the LIG-SEBS composite to enhance flexibility; (5) Laser Cut to shape the final electrode design; and (6) LIG Encapsulation, showing the finished electrode with LIG encased within the SEBS layer.

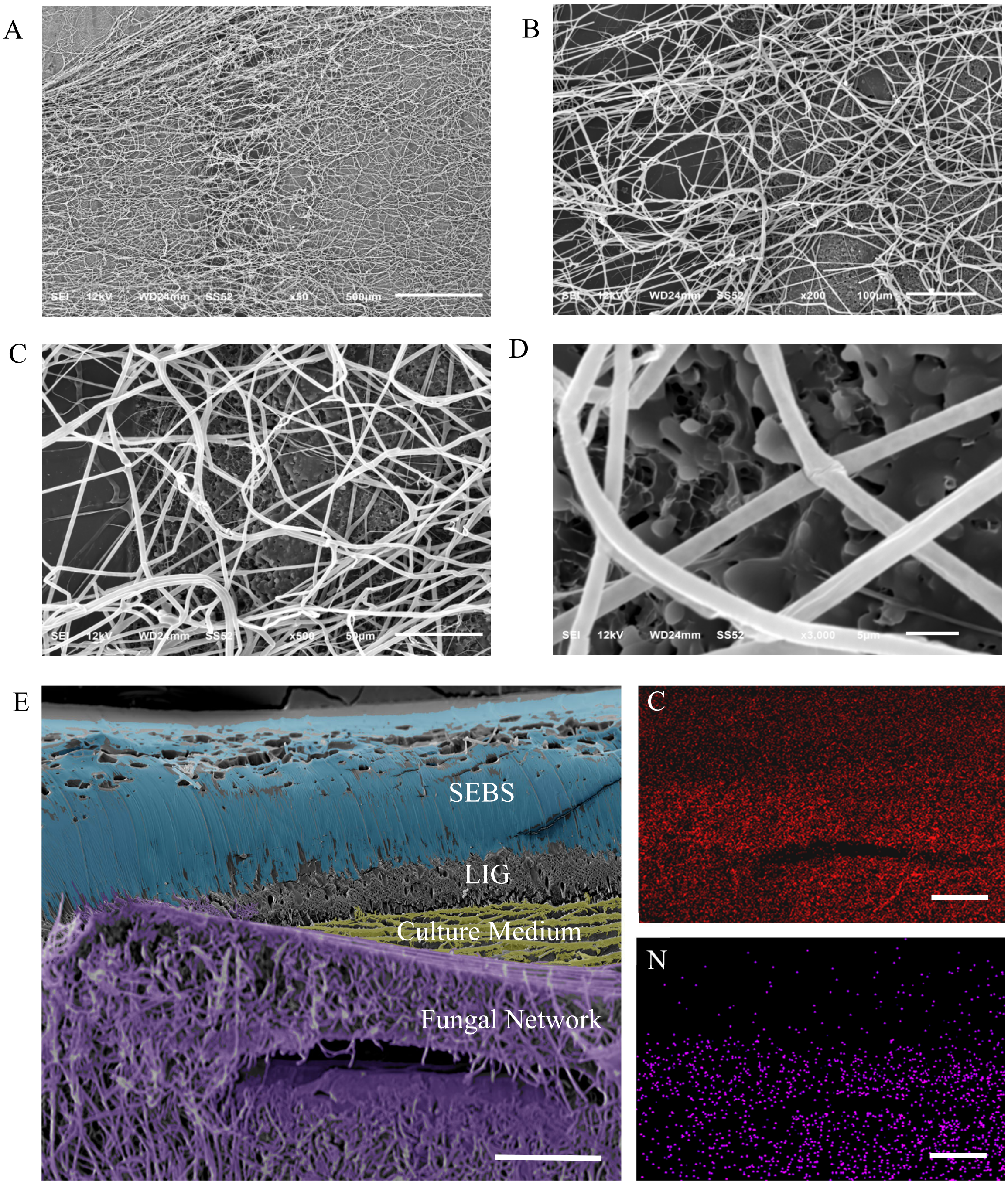


**Fig. 4. SEM-EDX images of *Benniella erionia* mycelium on LIG.** Mycelium on LIG electrodes (**A**) and higher-magnification views (**B**, **C** and **D**). SEM images demonstrate the interface between fungal mycelium and LIG electrodes in our Mycoelectronics. The micrographs reveal an intricate network of fungal hyphae spread across and closely adhering to the LIG surface at various magnifications. This extensive contact and integration between fungi and LIG, as visualized, indicates excellent connectivity between the biological and electronic components. **E,** The EDX image shown C, N elements are predominantly distributed in fungal regions (scale bar, 100 μm).

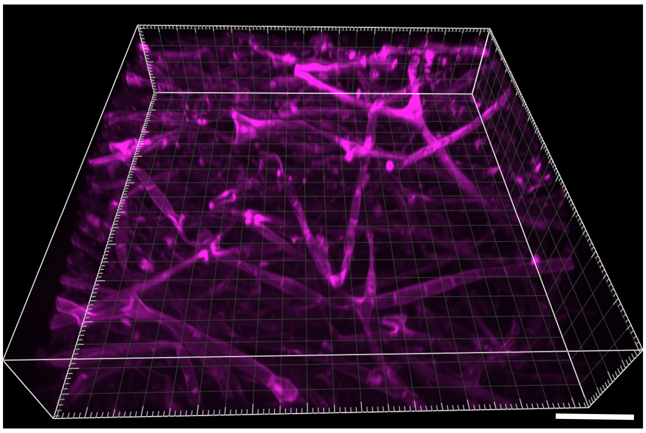

**Fig. S5. Confocal Imaging of Internal Hyphal Connections.** Confocal Z-stack image of *Benniella erionia* hyphae, revealing internal hyphal connections. The observed inter-hyphal linkages indicate potential continuous pathways for intracellular or intercellular transport (scale bar, 20 μm).

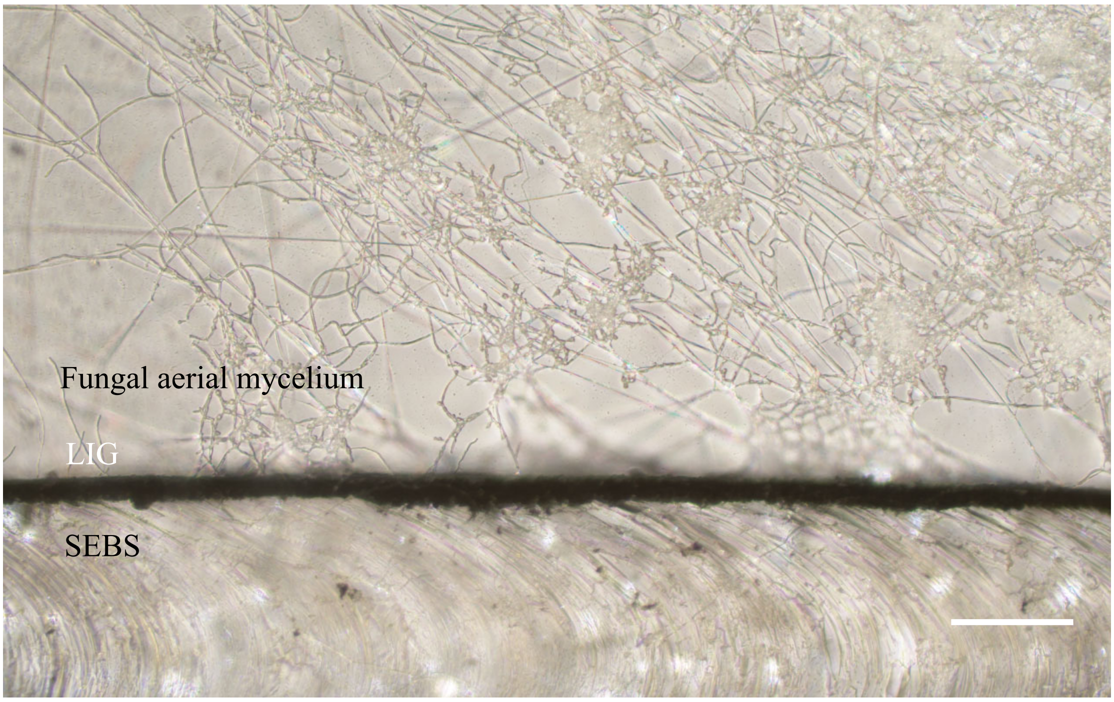


**Fig. S6. Optical photograph of the mycelium growing directly on the LIG electrode.** Fungal mycelium was directly co-cultured onto the LIG electrode without the addition of any medium. Cultivation was maintained at a controlled temperature of 23 °C and relative humidity of 30%. After 3 days of incubation, robust mycelial growth was observed, closely adhering to and spreading directly across the LIG surface. The numerous fine, branching structures visible are aerial hyphae, demonstrating the excellent biocompatibility and direct interaction between the fungus and the LIG substrate (scale bar, 100 μm).

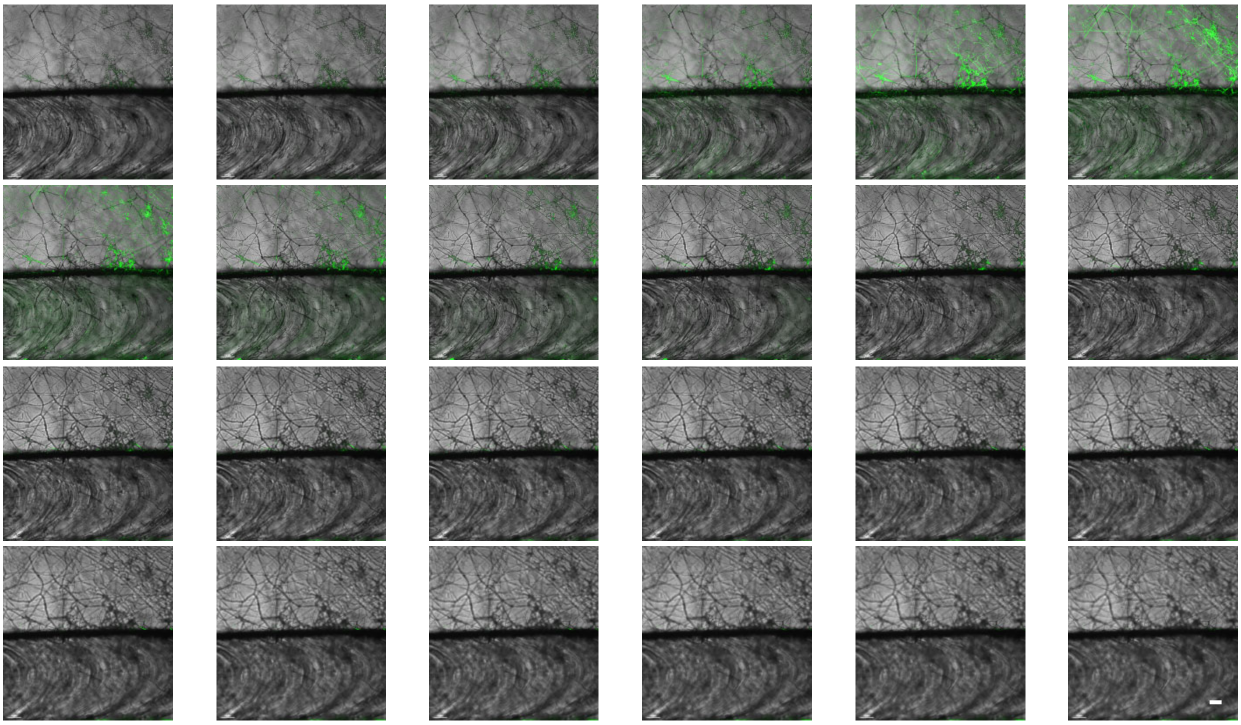

**Fig. S7. Cross-section of the fungal grow on LIG.** This series of confocal microscopy images presents a Z-stack of fungal mycelium growing on a LIG electrode with SEBS substrate. The dark horizontal line in each image represents the LIG-SEBS interface. Green fluorescence indicates fungal hyphae, utilizing the fungi's autofluorescence. As the focal plane moves through the Z-axis, we observe mycelial growth primarily on the LIG surface, with some hyphae extending above and below the LIG-SEBS layers. This spatial distribution of fungal colonization reveals how the mycelium integrates with our LIG-SEBS electrode system in various planes (scale bar, 100 μm).

A B

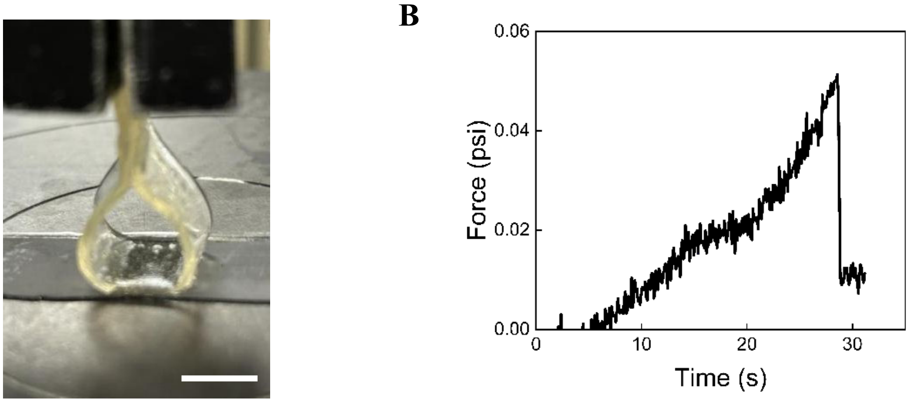


**Fig. S8. Force-time relationship in a loop tack test of a Gelatin based medium with LIG electrode.** (**A**) An optical image of the gelatin-based culture medium adhering to the LIG electrode during the test (scale bar, 1 cm). (**B)**, Force (in psi) as a function of time (in second). The adhesive force gradually increases to a maximum of approximately 0.05 psi over 30 seconds before rapidly decreasing, indicating the separation of the gelatin medium from the electrode. This behavior demonstrates the initial strong adhesion and subsequent controlled release between the gelatin-based medium and the LIG electrode, suggesting potential applications in stretchable Mycoelectronics. Corresponding video is provided in **Movie S2**.

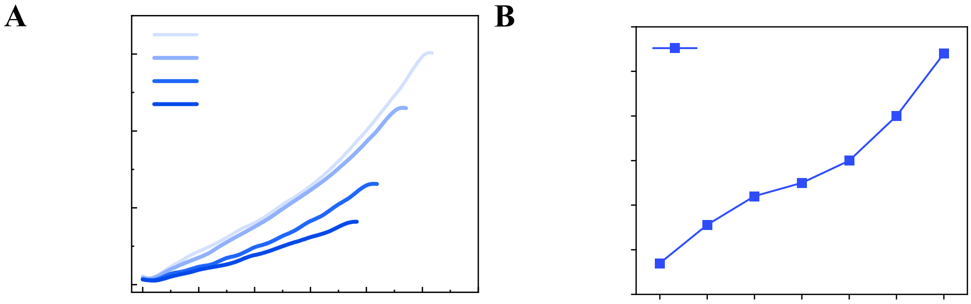


**Fig. S9. Culture medium mechanical and ion conductivity results.** (**A**) Strain-stress graph depicts the performance of different gelatin medium ratios (gelatin to 2% ME medium). It shows that higher gelatin content improves tensile performance. (**B**) the relationship between the growth of mycelium content in the medium and the change in conductivity of the medium/mycelium composite.

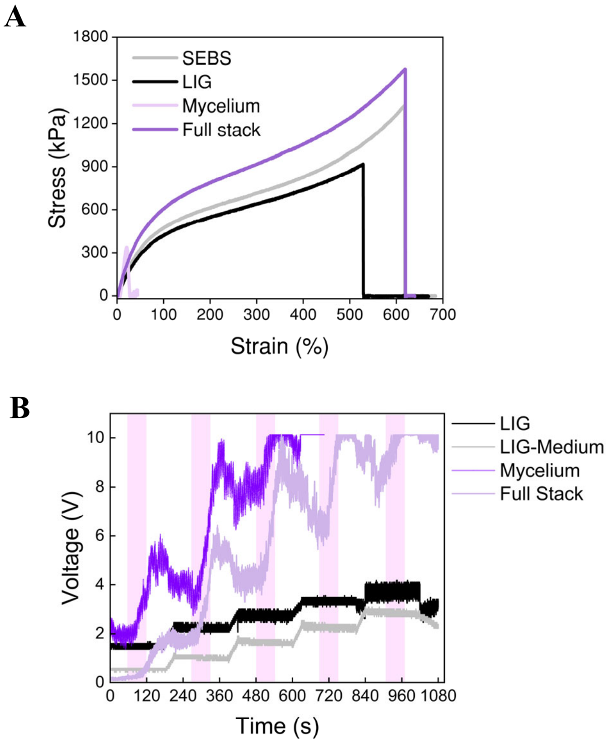


**Fig. S10. Stress strain test of each layer and entire Mycoelectronics.** (**A**) Stress-strain curves for SEBS, LIG, *Benniella erionia* Mycelium, and the full stack (Mycelium+medium+LIG+SEBS), with strain (%) on the x-axis and stress (Pa) on the y-axis. (**B**)Voltage response (1 nA constant current applied via PalmSens) and strain over time for LIG, LIG–SEBS, mycelium, and the full stack. The y-axis shows voltage (V) and the x-axis represents time (s). Pink shaded regions in the background denote periods of 45 °C thermal stimulation within each experimental cycle. Each cycle consisted of: 60 s rest → 60 s heating at 45 °C → 60 s rest → 30 s stretching to 10% strain → 60 s rest, repeated sequentially.

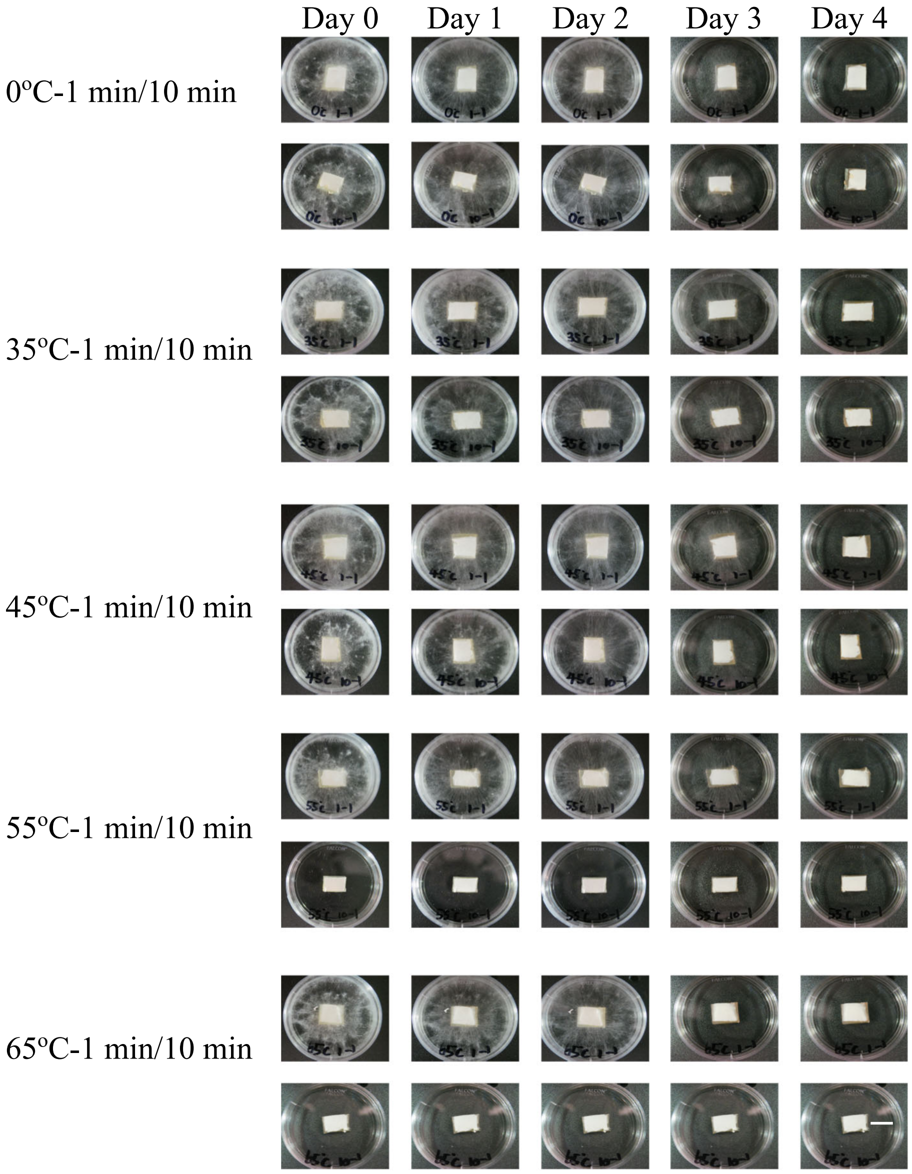


**Fig. S11. Temperature tolerance evaluation of *Benniella erionia* under varied thermal conditions.** Time-lapse imaging of fungal growth under different temperature exposures (0 °C to 65 °C) with short-term (1 min) and long-term (10 min) heat stimulation durations. Images were captured daily for 4 days following first day thermal treatment. Optimal growth was maintained between 0-45 °C, with notable growth inhibition observed at temperatures above 55 °C, particularly during extended exposure periods (scale bar, 1 cm).

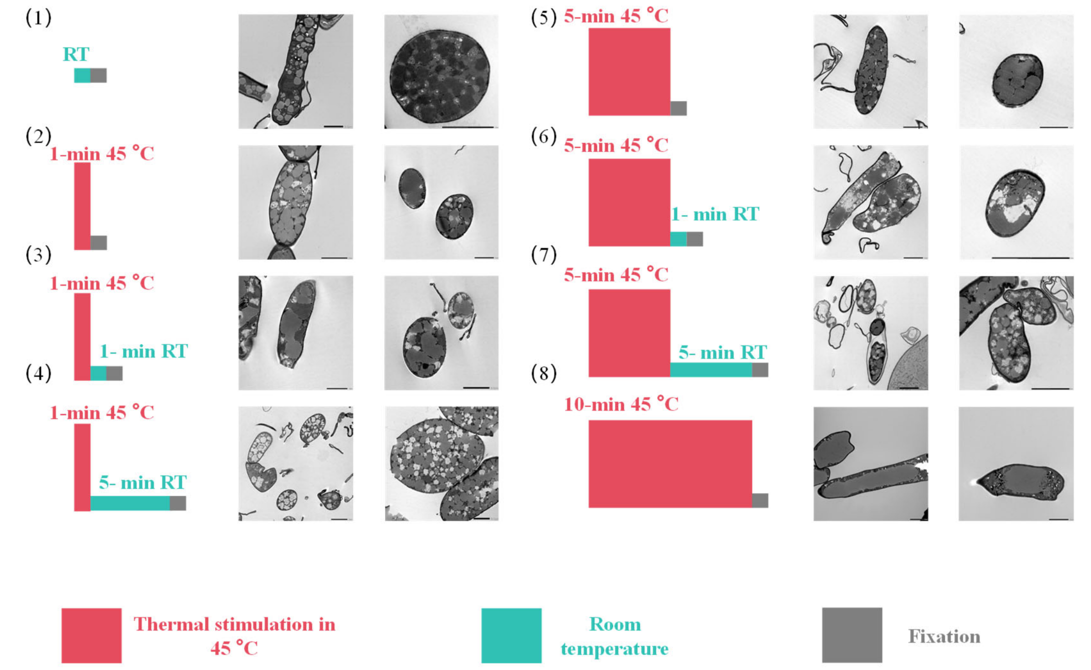


**Fig. S12. TEM micrographs of fungal cells subjected to different thermal treatments.** All hyphae were taken from the same Petri dish to minimize biological variability (scale bar, 2 μm). Eight treatment conditions were examined: (1) immediately fixed at room temperature (RT) without thermal stimulation; (2) 1-min thermal stimulation at 45 °C followed by immediate fixation; (3) 1-min thermal stimulation at 45 °C followed by 1-min rest at RT before fixation; (4) 1-min thermal stimulation at 45 °C followed by 5-min rest at RT before fixation; (5) 5-min thermal stimulation at 45 °C followed by immediate fixation; (6) 5-min thermal stimulation at 45 °C followed by 1-min rest at RT before fixation; (7) 5-min thermal stimulation at 45 °C followed by 5-min rest at RT before fixation; and (8) 10-min thermal stimulation at 45 °C followed by immediate fixation.

The images show that increased heat shock duration promotes vacuolar fusion, with the most extreme case (10-min thermal stimulation) producing a single large vacuole occupying almost the entire hyphal cross-section. Additionally, when hyphae were allowed to rest at RT after heat shock—regardless of whether the rest was 1 or 5 min—distinct intracellular voids appeared. These voids were pronounced after 5 min rest, suggesting that vacuoles formed during thermal stimulation shrink at lower temperatures, potentially due to osmotic pressure changes associated with temperature shifts. Together, these results indicate that thermal stress induces quick and reversible vacuolar remodeling in fungal hyphae, a phenomenon that can be detected in real time electrochemical (resistance) measurements, providing a mechanistic basis for the sensing function described in the main text.

A 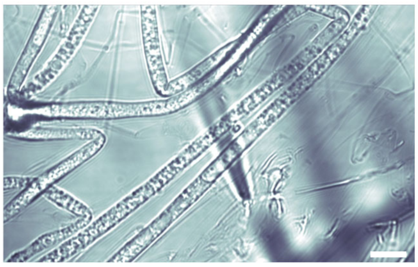 B 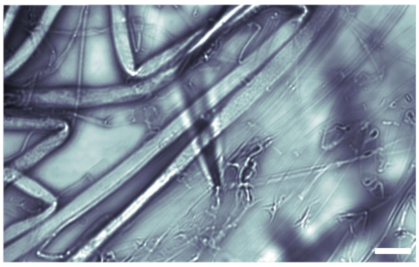

**Fig. S13. Bright-field imaging of fungal hyphae under thermal stimulation.** Bright-field images were captured using a Dragonfly confocal system with. (A) Baseline image before heating; (B) image after 35 °C thermal stimulation. The images reveal dynamic morphological changes of intracellular vacuoles, including volume change and fusion of vacuoles within single hyphae. These observations support findings from TEM and help confirm the thermally induced remodeling of fungal vacuolar architecture (scale bar, 5 μm). Corresponding video is provided in **Movie S6**.

A

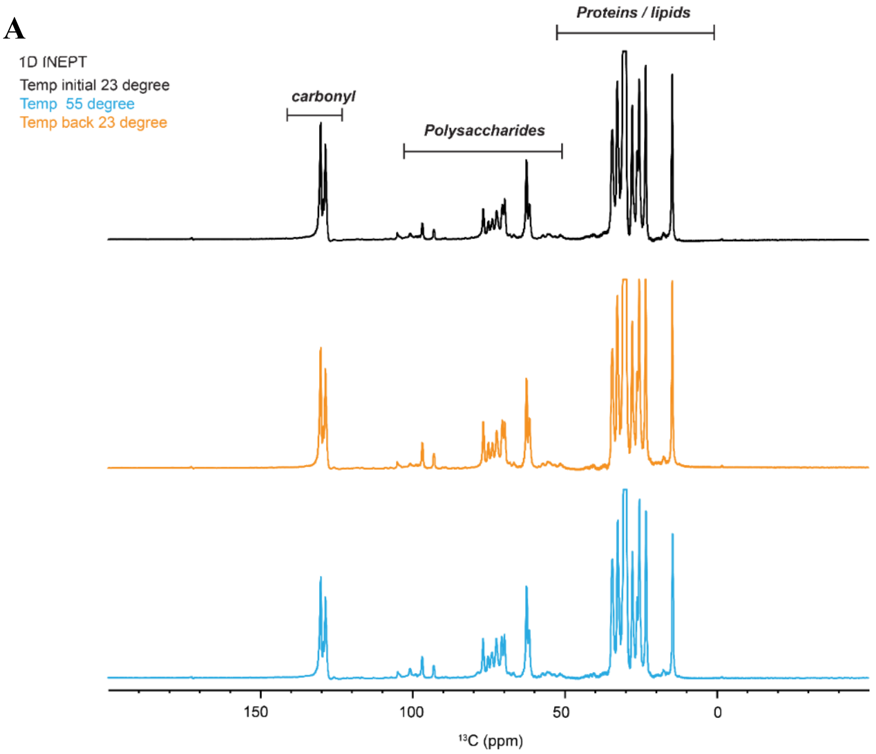

1D INEPT
Temp initial 23 degree
Temp 55 degree
Temp back 23 degree
carbonyl
Polysaccharides
Proteins / lipids
150
100
50
0
13C (ppm)


B

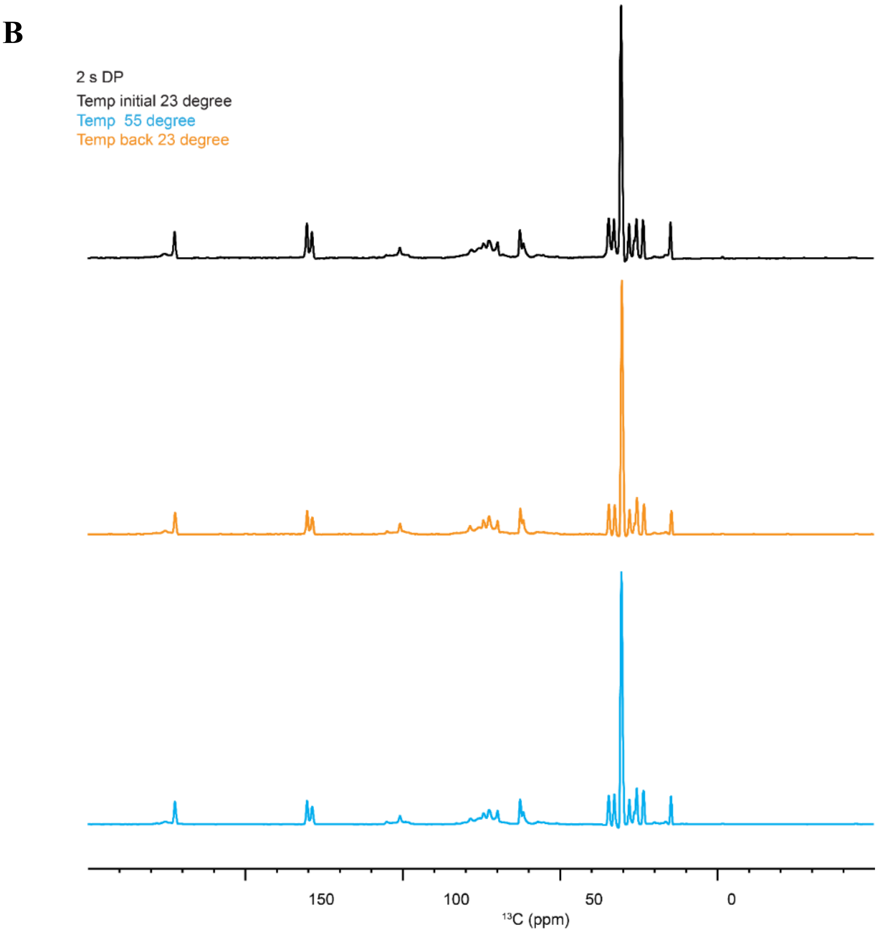

2 s DP
Temp initial 23 degree
Temp 55 degree
Temp back 23 degree
150
100
50
0
13C (ppm)

**Fig. S14. Solid state NMR result.** (**A**) 1D $^{13}$C refocused INEPT showing the different regions for carbohydrate and protein/lipid region. (**B**) 2 s DP spectra detecting the mobile molecules of fungal cell wall. It shows that there is not any significant change in the protein and lipid region after thermal stimulation. The one of $^{13}$C DP spectra measured with short recycle delay of 2 s that preferential detect mobile molecules has been acquired and it also shows that there is no effective change in the protein lipid region after the thermal stimulation. Overall, we can say that there is only change observed in the rigid part of the cell wall whereas mobile region remains same in both cases before thermal stimulation and after thermal stimulation.

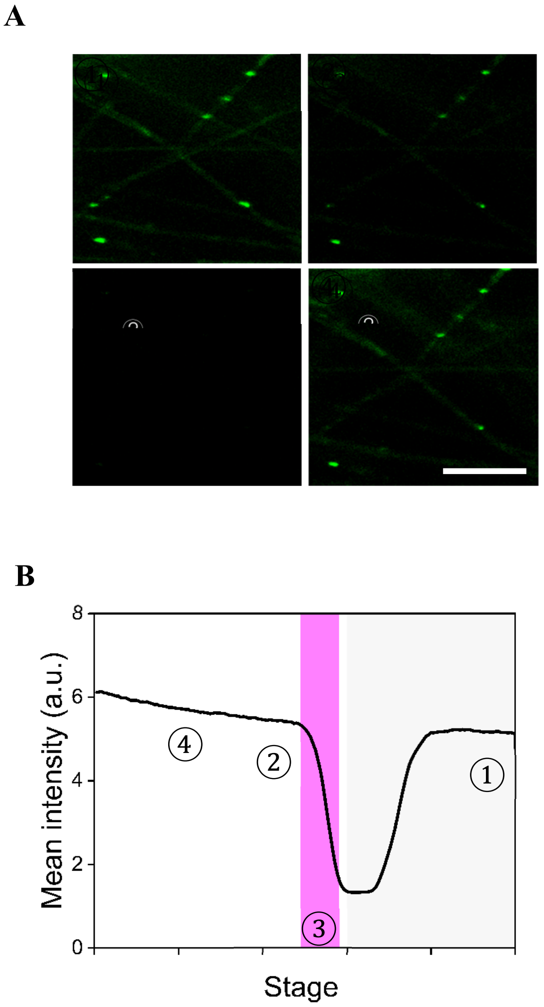


**Fig. S15. Fungal response under fluorescence microscope.** (**A**) Temperature-dependent fluorescence response of stained fungal mycelium under confocal microscopy (Scale bar, 25 μm). ① Initial state: The original fungal network structure and fluorescence intensity before thermal stimulation. ② Heating initiation: Thermal stimulation is applied. ③ Peak temperature: The fungal network at maximum thermal stimulation, just before the heat source is turned off. This image likely shows the most dramatic changes in network structure or activity. ④ Recovery state: The final image depicting the fungal network's state after cooling. (**B**) Fluorescence intensity changes were observed during heating and recovery. Corresponding video is provided in **Movie 7**.

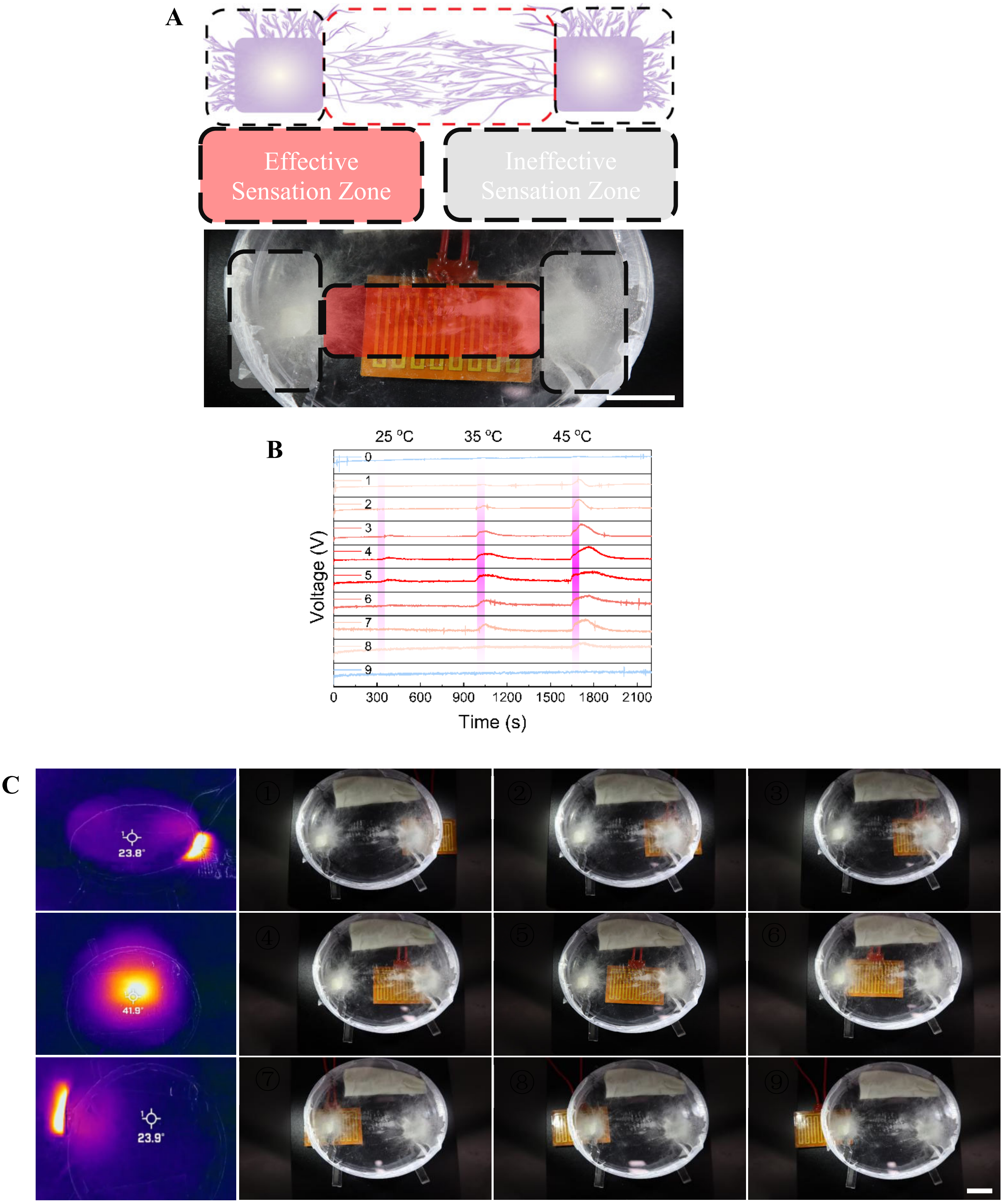


**Fig. S16. Definition of effective and ineffective temperature sensing regions in the Mycoelectronics.** (**A**) A schematic and an actual image of the experimental setup, depicting *Benniella erionia* mycelium growth across the electrode surface (scale bar, 1 cm). The central area, highlighted in red, represents the effective temperature sensing region, while the outer grey areas are ineffective. (**B**) graph of voltage responses over time for 9 testing points across the electrode, numbered 0 to 8. Three vertical pink bands indicate temperature stimulations at 25 °C, 35 °C, and 45 °C. The graph reveals that only the central testing points (3-5) exhibit significant voltage changes in response to temperature stimuli, corresponding to the effective sensing region defined in the left panel. Points 0-2 and 6-8 show minimal or no response, representing the ineffective sensing areas. **c,** the leftmost column shows thermal images from an IR camera, displaying temperature distributions at three key positions: heating pad to the right (23.8 °C), directly under (41.6 °C), and to the left (23.8 °C) of the Mycoelectronics. The 9 right images are photographs of the experimental setup, showing a transparent dome-like structure containing the fungal bioelectronic sensor and an orange heating pad in various positions (scale bar, 1 cm).

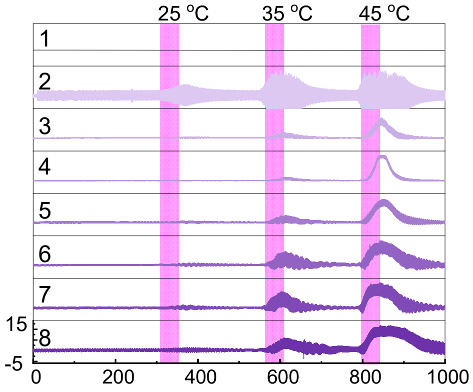


**Fig. S17. Voltage responses over eight days of testing.** Single channel Mycoelectronics achieved by inoculating two blocks of fungal on a pair of LIG electrodes after two days. Voltage fluctuations intensify was collected from Day 1 to Day 8, under different thermal stimulation with same duration. Despite the growing magnitude of responses, the pattern of each stimulus-induced reaction remains consistent across all days.

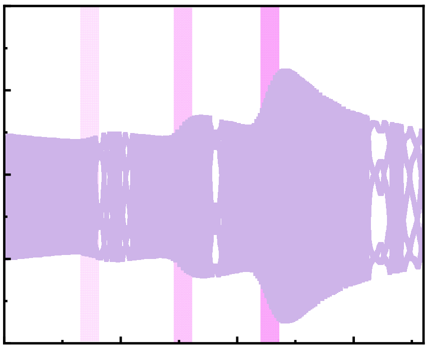

**Fig. S18. Long-term stability of Mycoelectronics stored for two months.** The recorded time–voltage curves at 25 °C, 35 °C, and 45 °C thermal stimulation exhibits consistent and proportional responses, confirming that the fungal mycelium remains viable and capable of reliable thermal sensing. This indicates that Mycoelectronics under well-controlled temperature and humidity, can be reactivated without any maintenance or nutrient replenishment.

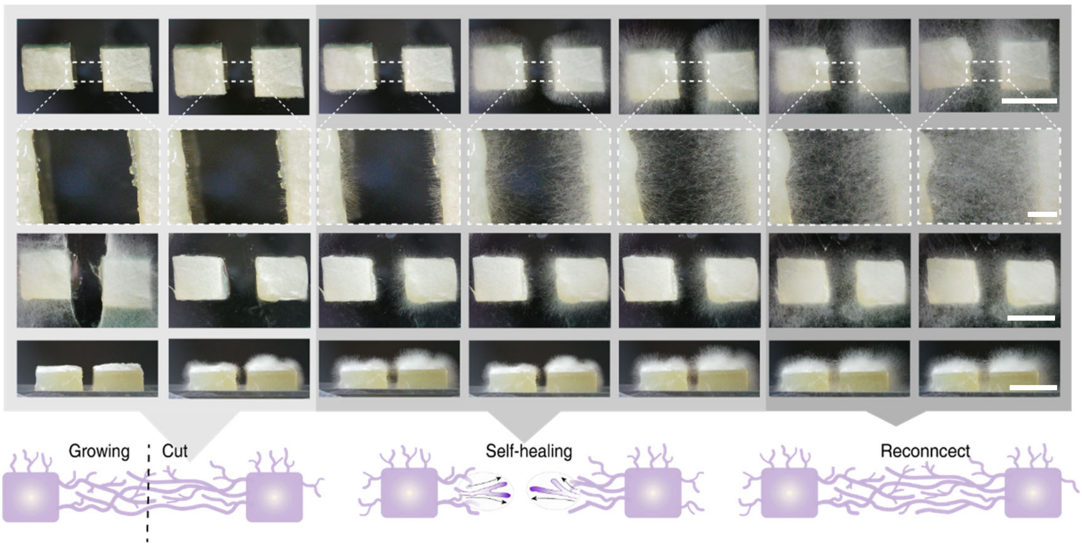


**Fig. S19. Self-healing capability of a single-channel Mycoelectronics.** The time-lapse sequence shows mycelial growth between inoculated with two *Benniella erionia* blocks, intentional disruption of the formed mycelial bridge, and subsequent rapid reconnection. Four rows of images capture the process: initial growth (scale bar, 1 cm), bridge formation (scale bar, 1 mm), cutting (scale bar, 1 cm), and self-healing (scale bar, 1 cm), aligns with the schematic at the bottom of the figure.

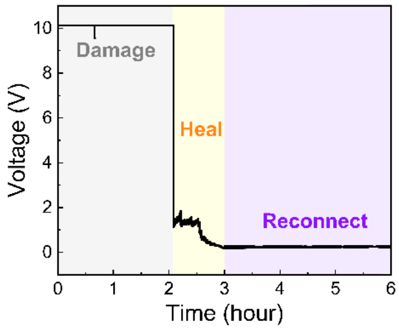


**Fig. S20. Continuous voltage monitoring demonstrating the self-healing process of the fungal network after mechanical damage.** Three distinct phases are observed: initial damage (gray region, 0-2 hours), healing phase (yellow region, 2-3 hours), and reconnection phase (purple region, 3-6 hours).

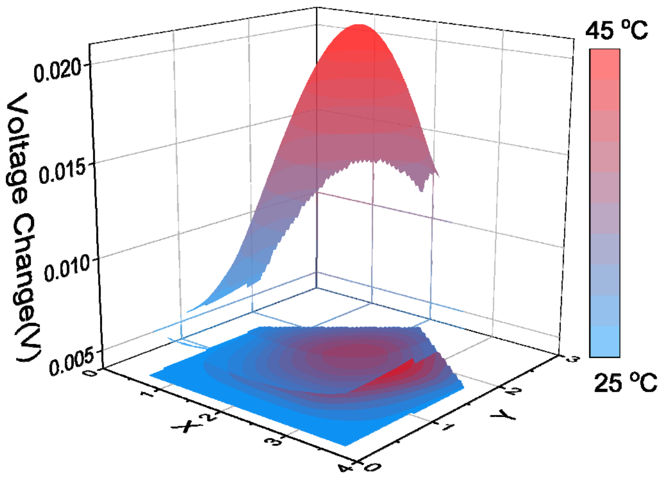


**Fig. S21. 16 channels mapping results.** Processed 3D temperature mapping result, cubic spline interpolation was implemented across the 16 nodes.

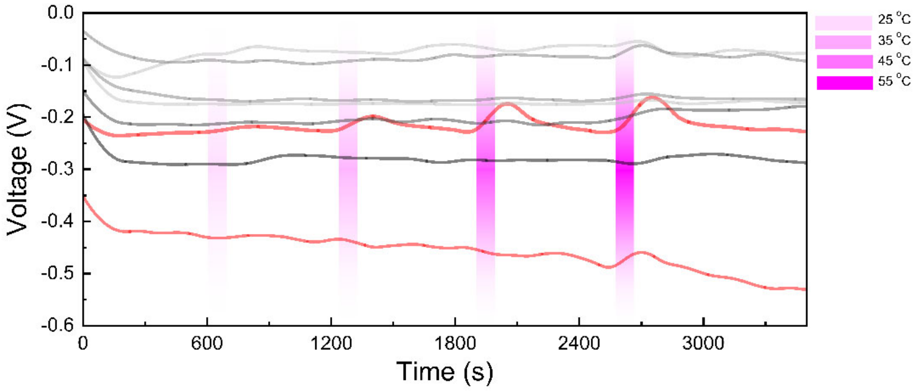


**Fig. S22. 3D electrode 8 channel results.** Time-dependent voltage responses recorded from a 3D electrode array during thermal stimulation. The multiple traces represent voltage measurements from electrodes at different distances from the heat source, with the red line indicating the electrode closest to the thermal stimulus. The voltage response amplitude correlates inversely with the distance from the heat source, demonstrating spatial sensitivity of the 3D electrode array. Gray and pink shaded regions indicate periods of thermal stimulation.

**Movie S1.** Fungal aerial mycelium growing toward and interacting with LIG electrodes
**Movie S2.** Loop tack test of gelatin based medium
**Movie S3.** Fracture dynamic of a freestanding fungal mycelium network
**Movie S4.** 3D bioprinting of bioink patterns on a LIG electrode array
**Movie S5.** Synchronized optical, infrared, and electrical data acquisition of Mycoelectronics thermoreceptor
**Movie S6.** .Real-Time observation of vacuolar fusion in hyphae
**Movie S7.** Voltage sensitive fluorescence response of mycelium to thermal stimulation
**Movie S8.** Rapid aerial mycelium network formation
**Movie S9.** Temperature-triggered smart window switch using a mycoelectronics thermoreceptor
**Movie S10.** Mycoelectronics activates cockroach leg motion
**Movie S11.** Mycoelectronics sensor activates robotic hand motion
